# Quantitative profiling of human translation initiation reveals regulatory elements that potently affect endogenous and therapeutically modified mRNAs

**DOI:** 10.1101/2024.02.28.582532

**Authors:** Cole J.T. Lewis, Li Xie, Shivani Bhandarkar, Danni Jin, Kyrillos S. Abdallah, Austin S. Draycott, Yixuan Chen, Carson C. Thoreen, Wendy V. Gilbert

## Abstract

mRNA therapeutics offer a potentially universal strategy for the efficient development and delivery of therapeutic proteins. Current mRNA vaccines include chemically modified nucleotides to reduce cellular immunogenicity. Here, we develop an efficient, high-throughput method to measure human translation initiation on therapeutically modified as well as endogenous RNAs. Using systems-level biochemistry, we quantify ribosome recruitment to tens of thousands of human 5′ untranslated regions and identify sequences that mediate 250-fold effects. We observe widespread effects of coding sequences on translation initiation and identify small regulatory elements of 3-6 nucleotides that are sufficient to potently affect translational output. Incorporation of N1-methylpseudouridine (m1Ψ) selectively enhances translation by specific 5′ UTRs that we demonstrate surpass those of current mRNA vaccines. Our approach is broadly applicable to dissect mechanisms of human translation initiation and engineer more potent therapeutic mRNAs.

**Highlights:**

- Measurement of >30,000 human 5′ UTRs reveals a 250-fold range of translation output
- Systematic mutagenesis demonstrates the causality of short (3-6nt) regulatory elements
- N1-methylpseudouridine alters translation initiation in a sequence-specific manner
- Optimal modified 5′ UTRs outperform those in the current class of mRNA vaccines

## Introduction

Genetic medicines are a potentially transformative class of therapeutics with diverse applications, including as vaccines, immunotherapies, and treatments for genetic disorders. Over the last few years, synthetic mRNAs have emerged as a front-runner among genetic medicine technologies, largely due to the overwhelming success of mRNA-based COVID-19 vaccines. mRNA-based therapeutics are relatively simple to design and produce, can rapidly induce therapeutic protein production, and act transiently without modifying cellular DNA. However, the broader use of mRNA therapeutics is currently limited by the amount of protein that can be produced.

Translation initiation is a rate-limiting process for protein synthesis in human cells. For most messages, translation initiation requires the concerted action of many eukaryotic initiation factors (eIFs) on an mRNA with a 5′ m^7^G cap^1^. The cap structure is recognized by a complex containing the cap-binding protein eIF4E, the DEAD-box RNA-dependent ATPase eIF4A, and the large scaffold protein eIF4G. In higher eukaryotes, eIF4G mediates ribosome recruitment by binding directly to the eIF3 subunit of 43S complexes that consist of a 40S small ribosomal subunit bound to eIF3, a ternary complex of eIF2•GTP•Met-tRNA_i_, and additional factors. The assembled complex scans the 5′ UTR from 5′ to 3′ to find the appropriate start codon, at which point the 60S large ribosomal subunit joins in a reaction requiring several additional factors. Increasing the rate of ribosome recruitment could substantially increase the potency of mRNA vaccines.

In endogenous mRNAs, translation-enhancing features are generally found in the 5′ untranslated region of mRNAs. Remarkably, changes in 5′ UTR sequences can vary the translation output of an mRNA more than 1,000-fold^2,3^. Some features that distinguish efficiently translated mRNAs include the presence of an m^7^G cap in an unstructured context at the 5′ end, an AUG initiation codon in a preferred Kozak sequence, shorter 5′ UTRs, lower GC content, and the absence of upstream initiation codons^4,5^. Beyond these minimal attributes, our understanding of the RNA-encoded elements that determine the translation output of mRNAs remains limited. Therefore, translational enhancers cannot yet be designed from first principles.

Naturally occurring translation enhancers may not function in therapeutic mRNAs, which include modified nucleotides to mask the RNA from immune sensors. Unmodified “native” mRNA triggers an intracellular innate immune response through RNA-surveillance mechanisms, including toll-like receptors (TLRs), the pattern-recognition receptor RIG-I, and the RNA-dependent kinase PKR^6–9^. Incorporation of modified nucleotides (e.g., N-1-methylpseudouridine) suppresses recognition by TLRs^10–12^. However, modified nucleotides affect RNA-RNA and RNA-protein interactions^13–15^ and are therefore likely to disrupt elements that normally promote translation in unmodified mRNAs.

Here we present <u>D</u>irect <u>A</u>nalysis of <u>R</u>ibosome <u>T</u>argeting (DART) as a facile approach to quantify translation initiation by more than 35,000 modified 5′ UTRs in human cell extracts. We find that human 5′ UTR-specific ribosome recruitment activity spans over 250-fold and is mediated by novel sequence-dependent mechanisms. Remarkably, the presence of N1-methypseudouridine affects ribosome recruitment to specific 5′ UTRs by more than 30-fold. DART measurements of ribosome recruitment directly predict protein synthesis from full-length mRNAs, with top-scoring 5′ UTRs outperforming those in the current class of mRNA vaccines. Our results uncover novel regulatory elements that potently affect translation initiation by human 5′ UTRs and establish DART as a powerful approach to engineer optimal 5′ UTRs for therapeutic mRNAs.

## Design

Facile, isoform-aware methods to study human translation initiation are currently lacking. Ribosome footprint-based approaches (Ribo-seq^16^, TCP-seq^17^, 40S footprinting^18^, etc.) are powerful methods to quantify ribosome occupancy at the gene level but lack information on the specific transcript isoforms and 5’ UTR sequences that recruited the ribosome. Isoform-specific polysome profiling approaches (TrIP-seq^2^, TL-seq^19^, etc.) are labor-intensive, require large amounts of input material, and do not decouple translation initiation from elongation and mRNA decay. DART, previously developed in budding yeast^3^, appeared to be a promising method to overcome these limitations. However, the original DART protocol required prohibitively large volumes of cell extract and was unable to test multiple conditions in a single experiment. We optimized the DART protocol for use in mammalian systems, reducing hands-on time and decreasing the required input material by two orders of magnitude. We incorporated a barcode-based multiplexing strategy and demonstrated its use by testing different RNA modifications within the same translation reaction and combining multiple reaction conditions onto the same sucrose gradient. These advances massively increase the throughput of DART, enabling researchers to measure the impact of a wide array of genetic, pharmacological, and RNA chemical perturbations on translation initiation. The use of a designed pool of 5′ UTRs in DART further allows direct testing of putative regulatory elements, moving beyond correlation to determine causality.

## Results

### Development of DART to quantify human 5′ UTR-mediated translational control

Translation initiation culminates in the recruitment of an 80S ribosome positioned at the start codon to begin polypeptide synthesis. 5′ UTRs play a critical role in this process, acting as a platform for binding of initiation factors necessary for ribosome recruitment. However, the features of 5′ UTRs that are responsible for conferring efficient initiation remain largely unknown, and the effects of modified nucleosides (e.g., N1-methylpseudouridine) are currently impossible to predict. We therefore sought to develop a high-throughput method to quantify and dissect the effects of 5′ UTR sequences and modifications on translation initiation. Direct Analysis of Ribosome Targeting (DART), which was recently described for measuring synthetic 5′ UTR activity in cell-free translation extracts from budding yeast^3^, appeared to be a promising strategy compatible with testing modified RNAs.

To adapt DART for use in a human system, we began by designing DNA oligonucleotide libraries containing over 35,000 Ensembl-annotated^20^ full-length human 5′ UTRs from 14,544 genes. The 5′ UTR sequences range from 10-230 nucleotides in length followed by at least 27 nucleotides of coding sequence to provide a binding site for the initiating ribosome. Upstream AUGs were removed for simplicity. Oligos include a T7 promoter at the 5′ end for in vitro transcription and a common primer binding site at the 3’ end for library construction. We transcribed the DNA library and enzymatically added a 5′ methylguanosine cap to produce an RNA pool that reflects endogenous mRNA 5′ ends. RNA pools were incubated in an in vitro translation reaction with Hela cytoplasmic lysate, which recapitulates cap-stimulated translation over a wide range of mRNA concentrations (**Figure S1A and B**). Translation reactions contained cycloheximide to stabilize recruited ribosomes during sucrose gradient centrifugation, which separated ribosome-bound RNAs from those that failed to recruit a ribosome. Following RNA recovery from the 80S fraction, we prepared Illumina sequencing libraries and calculated a ribosome recruitment score (RRS) as the relative abundance of 80S-bound RNA compared to an input control library (**Figure 1A**).

**Figure 1.**
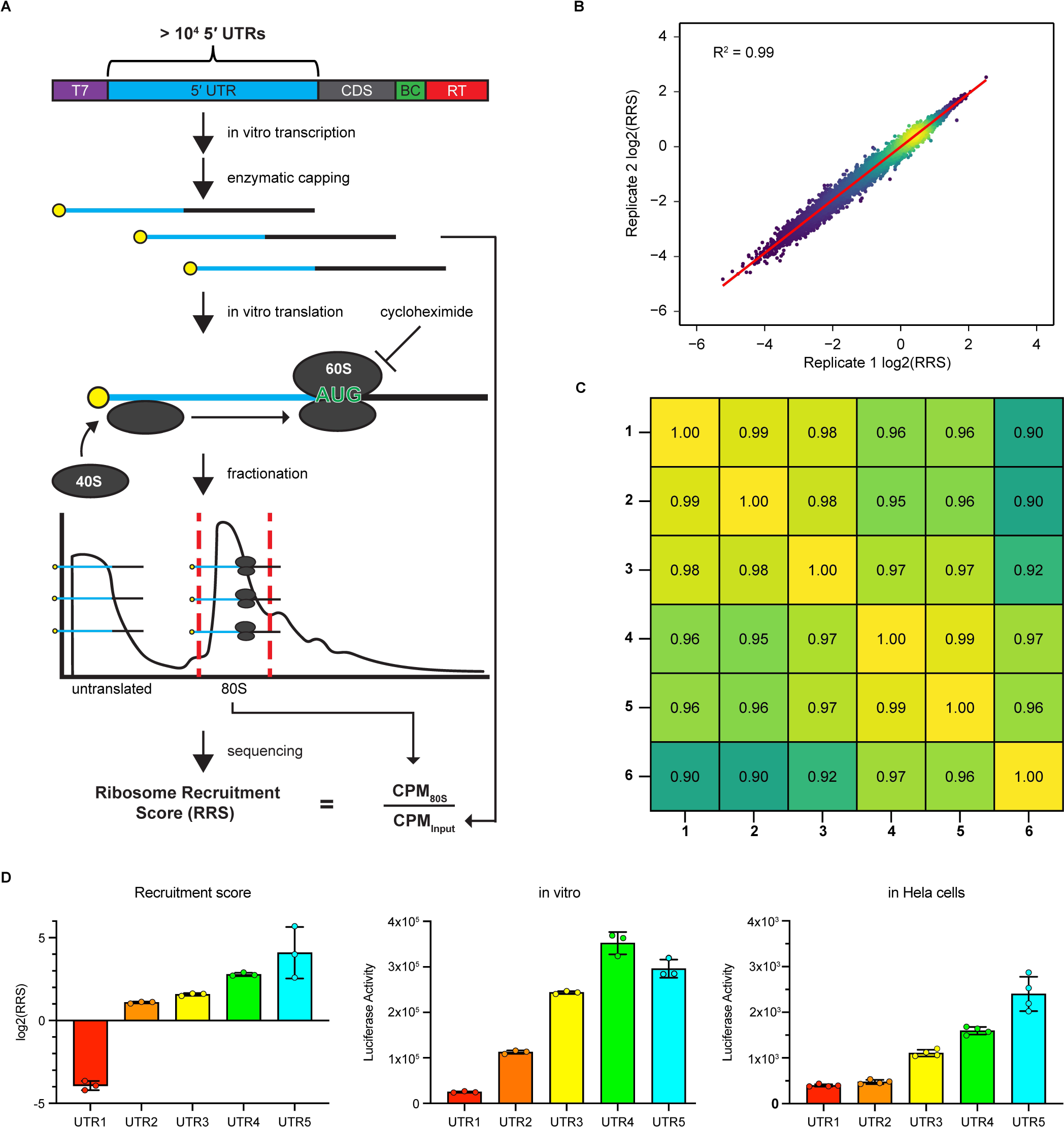
DART quantifies human 5′ UTR-mediated translational control over a 250-fold range. A) Schematic of the DART workflow. The sequences in the DNA pool contain a T7 promoter followed by > 27nt of coding sequence and a reverse transcriptase binding site for library preparation. Endogenous 5′ UTR sequences were derived from Ensembl annotations. B) Human DART reproducibly measures ribosome recruitment over a 250-fold range. C) Comparison of six DART replicates. D) Ribosome recruitment scores predict 5′ UTR-driven translational activity in full-length firefly luciferase reporter mRNAs *in vitro* (HeLa cytoplasmic lysate, middle) and in cells (HeLa cell transfection, right).

Human DART reproducibly quantified 5′ UTR activity spanning over a 250-fold range (R^2^ = 0.90-0.99, **Figure 1B and C**), highlighting the extensive translational control exerted by human 5′ UTRs. We selected 5′ UTRs that spanned the range of ribosome recruitment scores (**Figure 1D**, left panel) and cloned them upstream of the firefly luciferase (fluc) coding sequence. These reporters were in vitro transcribed, m^7^G capped, and translated in Hela cytoplasmic extract and via transfection into Hela cells. We observed good agreement between ribosome recruitment scores from the DART assay and the amount of protein synthesized (**Figure 1D**). Thus, human DART enables high-throughput quantification of 5′ UTR activity, and the DART measurements predict protein synthesis in the context of full-length mRNAs.

### Systematic testing shows repression by C-rich sequence motifs

We sought to determine 5′ UTR sequence elements that could explain the observed differences in ribosome recruitment. We selected the 100 most active and 100 least active 5′ UTRs from our initial DART analysis and generated a new library in which we systematically deleted 6-nucleotide segments scanning along the full length of each 5′ UTR (**Figure 2A**). DART analysis on this library of over 6,000 variants of the initial 200 sequences identified hundreds of putative translational enhancer and repressor elements (**Figure 2B**). An example of a putative translational enhancer within the 5′ UTR of TMSB15B is shown in **Figure 2C**, where deletion of cap-proximal nucleotides reduced ribosome recruitment more than fourfold.

**Figure 2.**
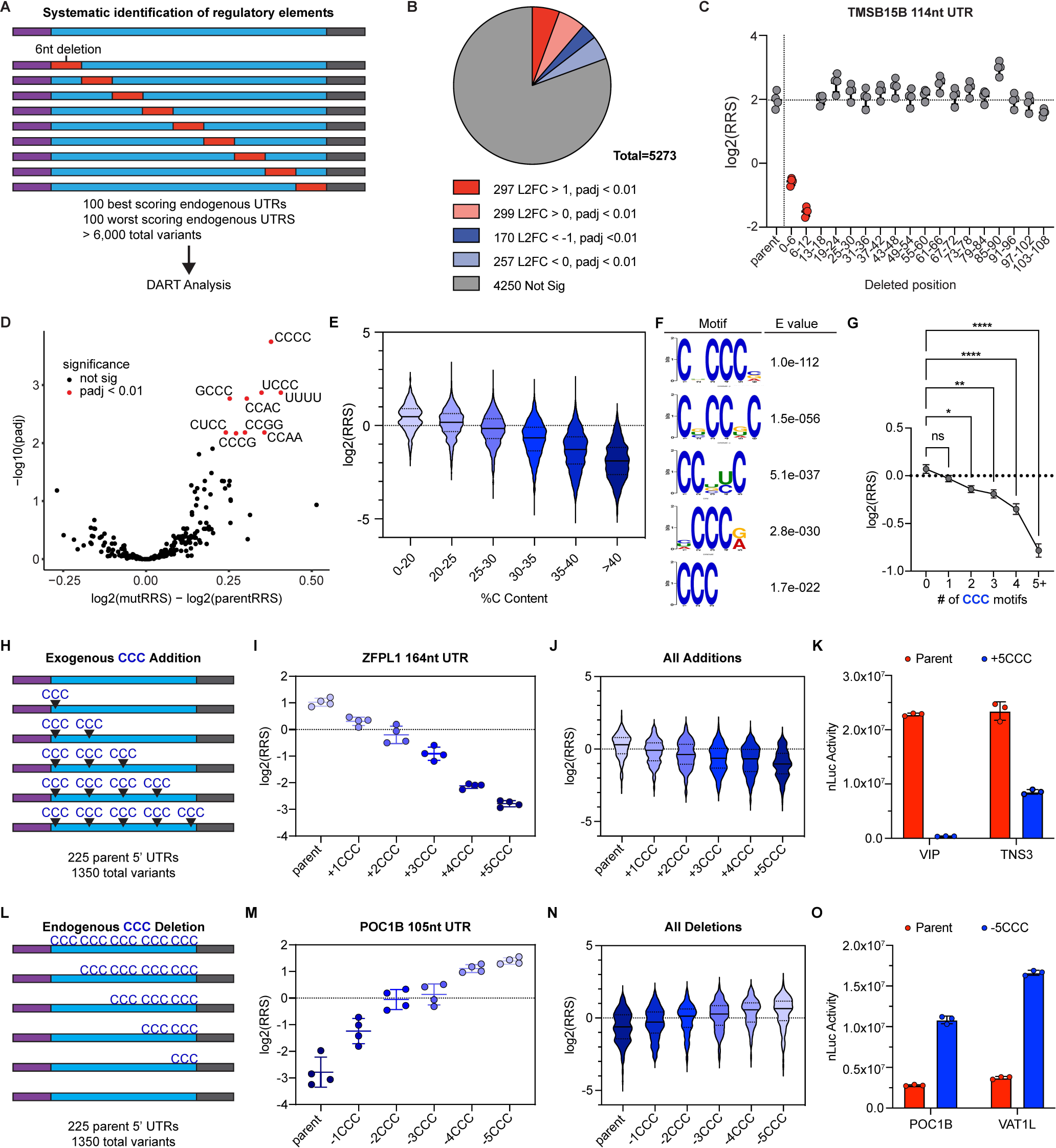
CCC motifs repress Vaccinia-capped 5′ UTR activity. A) Scanning deletion library design for systematic identification of regulatory elements. B) Scanning deletion analysis identifies hundreds of hexamers that significantly increase (red) or decrease (blue) ribosome recruitment (Benjamini-Hochberg adjusted p < 0.01). C) Translational enhancer in the TMSB15B 114 nucleotide 5′ UTR. Deletion of nucleotides 1-6 or 7-12 (red) reduces ribosome recruitment by over 4-fold. D) Deletion of C-rich elements increases ribosome recruitment. Volcano plot of tetramer sequences deleted at least 30 times covered in the scanning deletion library. Tetramer sequences that significantly altered ribosome recruitment are highlighted in red (Benjamini-Hochberg adjusted p < 0.01). E) Global trend of reduced ribosome recruitment with increasing cytidine content. 5′ UTRs binned by percent cytidine. F) Enrichment of C-rich sequence motifs in bottom 10% of 5′ UTRs by RRS. E values for motif enrichment determined by DREME. G) The number of CCC trinucleotide motifs correlates with decreased ribosome recruitment. Analysis of 2295 5′ UTRs with 30-35% cytidine overall, binned by the number of CCC motifs they contain. The data are represented as mean log2(RRS) for each bin (error bars = 95% CI) *p < 0.05, **p < 0.01, ***p < 0.001, ****p <0.0001 by one-way ANOVA and Tukey’s multiple comparisons test. H) Library design to test CCC motif dose-dependence using exogenous CCC additions. I) and J) CCC motifs repress ribosome recruitment in a dose-dependent manner. CCC sequences were iteratively added to 5′ UTRs and these UTRs were tested by DART. Repressive effect for ZFPL1 (J) and across all 5′ UTRs tested (K, n = 225 parent 5′ UTRs, 1350 total variants). K) 5′ UTR CCC motifs repress translation of luciferase mRNAs. L) Library design to test CCC motif dose-dependence by deletion of endogenous CCC motifs. M) and N) Removal of CCC motifs from human 5′ UTRs increases ribosome recruitment. CCC motifs were iteratively deleted from 5′ UTRs and tested by DART. The effect of successive CCC removal is demonstrated for the 5′ UTR of POC1B (M) and across all UTRs tested (N, n = 225 parent 5′ UTRs, 1350 total variants). O) Deletion of endogenous CCC motifs increases the translation of mRNAs.

To identify common translational regulatory motifs, we quantified the impact of tetramer sequences on ribosome recruitment. Focusing on sequences that were tested at least 30 times across the scanning deletion library, we observed that the removal of C-rich elements significantly enhanced ribosome recruitment (**Figure 2D**). Accordingly, we noted a striking anticorrelation between cytosine content and ribosome recruitment (**Figure 2E**) that was not observed for other nucleotides (**Figure S2A**). On average, 5′ UTRs with C content below 20% were 5.6-fold more active than 5′ UTRs with greater than 40% C. These results were further corroborated by an unbiased search for enriched motifs within the worst-performing 5′ UTRs. DREME motif enrichment analysis^21^ (Methods) of 5′ UTRs in the lowest decile of ribosome recruitment activities identified C-rich motifs (**Figure 2F**). Interestingly, 3 out of the top 5 enriched motifs contained a CCC trinucleotide element. To determine if CCC motifs affected ribosome recruitment beyond the overall C content of the 5′ UTRs, we examined 5′ UTRs containing 30-35% C (n = 2,295 UTRs). Within this group of 5′ UTRs, more CCC motifs correlated with less ribosome recruitment (**Figure 2G**).

Our analysis suggested that CCC elements within 5′ UTRs depress the translational activity of synthetic mRNAs. To test this directly, we generated a pool of 5′ UTR sequences in which up to 5 CCC motifs were iteratively added to 225 endogenous 5′ UTRs and performed DART on these sequences (**Figure 2H**). We observed a significant and dose-dependent decrease in ribosome recruitment to these 5′ UTRs upon addition of CCC elements (**Figures 2I and J**). We further validated the effect of CCC motifs in repressing translational output of full-length luciferase mRNAs (**Figure 2K**).

Given that the addition of exogenous CCC motifs was sufficient to repress translation, we tested whether the removal of endogenous CCC elements would conversely increase translation. Using a similar strategy to the CCC additions, we iteratively deleted up to 5 naturally occurring CCC elements within 225 5′ UTRs and performed DART on these sequences (**Figure 2L**). Sequential deletion of CCC elements resulted in a dose-dependent increase in ribosome recruitment to these 5′ UTRs (**Figures 2M and N**) and increased protein synthesis in full-length luciferase mRNAs (**Figure 2O**). These data demonstrate the strength of DART to rapidly iterate through 5’ UTR pools, from unbiased systematic discovery to designed variants that move from correlation to causation.

### Pervasive effects of RNA sequence and structure on enzymatic capping

CCC motifs have the potential to base-pair with the GGG present at the 5′ end of these RNAs as a part of the optimal T7 RNA polymerase promoter. Vaccinia capping enzyme (VCE) is known to require the 5′ end to be accessible to install an m^7^G cap on an RNA substrate^22^. As the addition of a cap increased translation activity by ∼40-fold in these extracts, variable capping efficiency is likely to significantly impact ribosome recruitment scores. We therefore hypothesized that increased numbers of CCC motifs within 5′ UTRs decreased ribosome recruitment by reducing VCE capping efficiency by sequestering the 5′ ends into inaccessible secondary structures. To determine the secondary structure of the 5′ UTR RNA library, we performed chemical probing with dimethyl sulfate (DMS-MaPseq^23^). Briefly, 5′ UTR pools were folded *in vitro*, treated with DMS to probe unpaired A and C residues, and reverse transcribed and sequenced. As expected, DMS treatment specifically increased mutation rates at A and C, and RNA folding led to significant protection compared to denatured controls (**Figure S2B,C**). DMS reactivity was used to constrain 5′ UTR folding *in silico* to determine the pairing probability for regions of interest (Methods, **Figure S2D**). Consistent with our hypothesis, 5′ UTRs with low activity exhibited significantly more pairing at their 5′ ends (**Figure S2E**).

To determine if the repressive effect of CCC motifs was due to inhibition of enzymatic RNA capping, we compared luciferase reporters with the same 5′ UTR sequences capped two ways, enzymatically (VCE) and co-transcriptionally (CleanCap AG). The effects of adding or deleting CCC motifs were substantially blunted with CleanCap (compare **Figures S2F and S2G** to **Figures 2L and 2O**). In further support of a repressive effect of CCC elements on VCE capping efficiency, the addition 5 CCC elements significantly reduced the stimulatory effect of VCE on protein production (**Figure S2H**) compared to parental sequences. Conversely, the deletion of 5 CCC elements significantly enhanced the stimulatory effect of capping with VCE (**Figure S2I**). Together, these results establish a general inhibitory effect of short CCC motifs on VCE-capped mRNAs, which has implications for research and therapeutic mRNA design.

### Familiar 5′ UTR determinants exert modest effects on ribosome recruitment with many exceptions

To determine features of 5′ UTRs that directly impact translation initiation on capped mRNAs, we generated a new library of 51,596 co-transcriptionally capped 5′ UTRs using CleanCap AG, which generates >94% capped RNA^24,25^ and performed DART analysis. We observed a weak correlation between the ribosome recruitment activity of VCE-capped versus co-transcriptionally capped 5′ UTRs compared to replicate reproducibility (R^2^ = 0.17 versus 0.96 - 0.83, **Figure S3A**), indicating VCE sequence bias causes pervasive effects. We sought to determine if the observed negative effect of cap-proximal structure formation **(Figure S2E)** was solely due to inhibition of capping by VCE. DMS-MaPseq profiling of the co-transcriptionally capped RNA library revealed a substantially reduced, but still significant impact of cap-proximal pairing on ribosome recruitment (**Figure S3B**). The remaining translational repression by cap-proximal RNA structure is consistent with reduced association with the cap-binding complex^26,27^.

We began our analysis of translational regulatory features within 5′ UTRs by examining general trends. We observed a significant anticorrelation of 5′ UTR length and ribosome recruitment score, with UTRs less than 80 nts having higher activity (**Figure 3A**). A minimum of 12-14 nts are needed to span the distance from the ribosomal P site to the cap^28,29^. It is therefore notable that UTRs less than 14 nts in length recruited ribosomes efficiently (**Figure 3B**) and promoted high levels of reporter protein synthesis (**Figure S3C**). RNA folding stability (**Figure 3C**), and GC content (**Figure 3D**) were negatively correlated with ribosome recruitment activity. However, many 5′ UTRs deviated from the global trends. Some highly structured 5′ UTRs were among the most efficient ribosome recruiters (**Figure S3D and S3E**).

**Figure 3.**
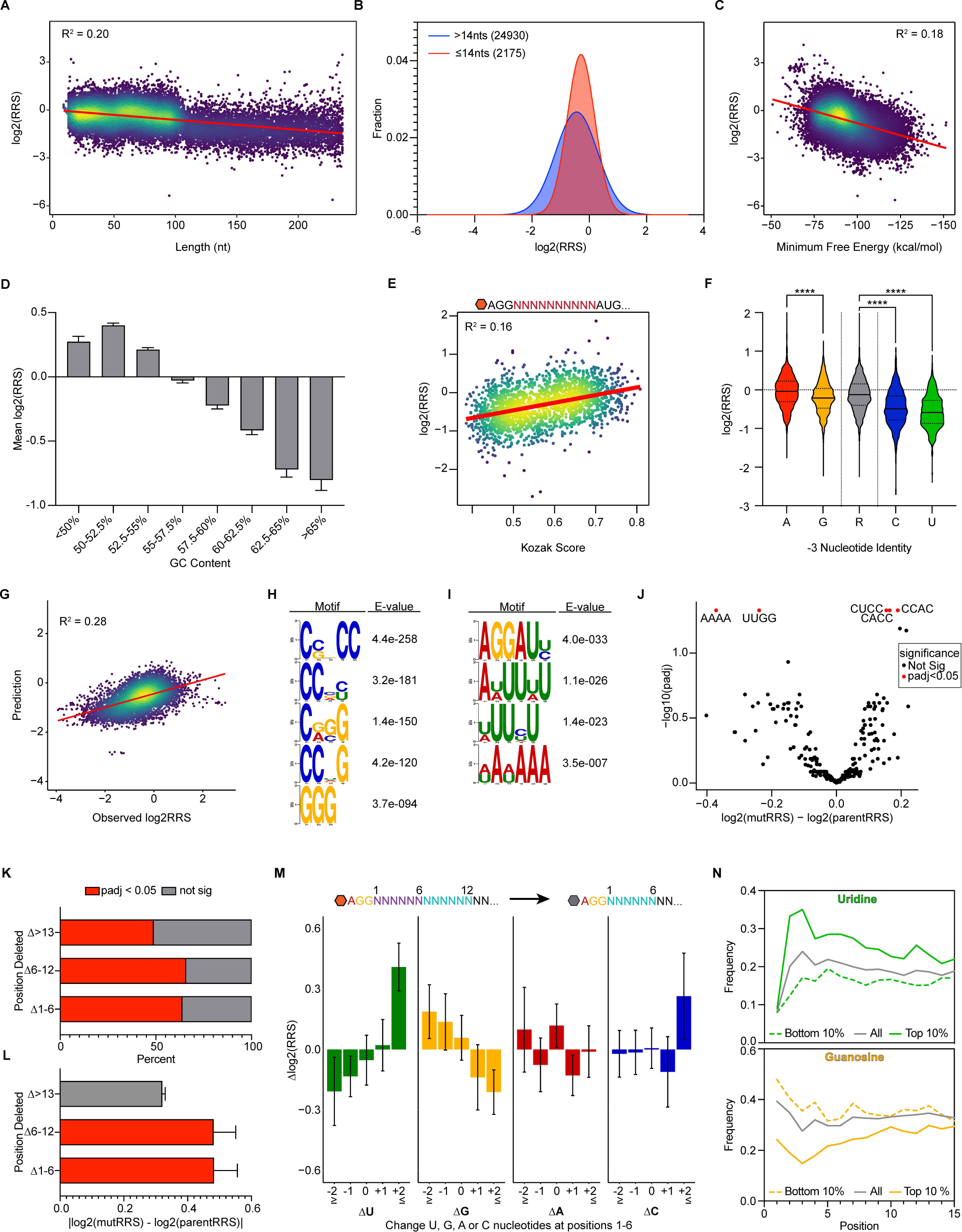
Known regulatory elements explain little of the observed variation in 5′ UTR-specific translation activity. A) Effect of UTR length on ribosome recruitment. B) Histogram of log2(RRS) scores for 5′ UTRs shorter than 15 nucleotides (red) or greater than 15 nucleotides (blue). C) Predicted minimum free energy negatively correlates with ribosome recruitment and explains 18% of the variability observed in DART. D) Impact of GC content on ribosome recruitment. Sequences were binned according to their percent GC. Data are represented as mean log2(RRS) for each bin (error bars = 95% CI). E) Kozak sequence strength promotes ribosome recruitment to random 10-nucleotide 5′ UTRs. UTRs scored based on conformity to the consensus human Kozak sequence and plotted against log2(RRS). F) Pyrimidines at the −3 position correlate with worse ribosome recruitment. 5′ UTRs were binned based on their nucleotide identity at −3 relative to the start codon. ****p < 0.0001, unpaired Welch’s t-test. G) Length, predicted minimum free energy, GC content, and Kozak strength are insufficient to determine 5′ UTR ribosome recruitment activity. A linear regression model incorporating these features explains 28% of the variance in DART. H) and I) Sequence motifs enriched in H) worst 10% and I) best 10% of 5′ UTRs by log2(RRS) from DART. J) Volcano plot of tetramer sequences deleted at least 30 times covered in the scanning deletion library. Tetramer sequences that significantly altered ribosome recruitment are highlighted in red. K and L) Deletions of cap-proximal nucleotides significantly affect ribosome recruitment. 5′ UTRs were binned by the position of deleted nucleotides. Deletions of nucleotides 1 through 6 or 7 through 12 K) were more likely to cause a significant change (Benjamini-Hochberg adjusted p < 0.01) in ribosome recruitment and L) caused larger magnitudes of change in ribosome recruitment by DART. M) Gain of uridines and loss of guanosine nucleotides within the first 6 nucleotides increases ribosome recruitment in a dose-dependent manner. 5′ UTRs from the scanning deletion library were binned based on the change in the number of each nucleotide. Change in RRS relative to the respective parent sequences is plotted on the y-axis. N) Uridines or guanosines are enriched in highly active or minimally active 5′ UTRs, respectively. UTRs were binned based on the top 10% (solid lines) or bottom 10% (dashed lines) by ribosome recruitment scores. Plot displays the percent of UTRs containing uridine (top) or guanosine (bottom) at each position.

As the 43S complex scans along the 5′ UTR, the nucleotides surrounding the AUG start codon, referred to as the Kozak sequence, play an important role in determining where translation will begin^30,31^. To quantify the impact of the Kozak sequence directly, we generated a library of 2000 random 10-nucleotide 5′ UTRs and measured their ribosome recruitment activity. We then scored each 10mer UTR based on their similarity to the consensus human Kozak sequence^32^ (GCCRCC<u>AUG</u>G, R = purine), and observed a positive correlation between Kozak strength and ribosome recruitment (**Figure 3E**). Consistent with recognition of a purine at the −3 position^31,33^, 10mers containing a U (n = 461) or C (n = 513) at this position recruited significantly fewer ribosomes than those with the consensus −3R (n = 1011) (**Figure 3F**). However, most expressed human 5′ UTRs contain a purine at −3 indicating that features other than Kozak strength are responsible for the wide range of ribosome recruitment observed (**Figure S3F and S3G**).

### Cap-proximal nucleotide composition affects ribosome recruitment

Most 5′ UTRs showed activities that were not explained by the general trends in length, structure, or Kozak score. A linear regression model incorporating these features was only able to predict 28% of the variability in DART (**Figure 3G**). To identify sequence elements outside the Kozak region that regulate translation, we performed motif enrichment analysis on the least and most active deciles of 5′ UTRs in our library. We observed C- and G-rich elements were enriched amongst the worst performing 5′ UTRs (**Figure 3H**), indicating that C-rich elements repress translation initiation outside of their negative effect on enzymatic capping. Amongst the top-performing UTRs, U/A-rich elements were enriched (**Figure 3I**), suggestive of an enhancing effect. To directly test the impact of short sequence elements on initiation, we repeated DART analysis on the systematic deletion library (**Figure 2A**) and noted that deletion of individual C-rich tetramers had a significant enhancing effect on ribosome recruitment (**Figure 3J**).

Deletions of cap-proximal nucleotides (Δ1-6 or Δ7-12) were more likely to significantly alter ribosome recruitment (**Figure 3K**) and caused a larger magnitude of change (**Figure 3L**) than deletions in the remainder of the UTR sequence (Δ>13). Deletion of the 5′-most 6 nucleotides results in replacement of the cap-adjacent nucleotides, allowing us to directly assess the impact of changing nucleotide composition in this region. We observed that gain of Us and loss of Gs at positions 1 through 6 significantly increased or decreased ribosome recruitment, respectively, in a dose-dependent manner (**Figure 3M**). Accordingly, across the entire 5′ UTR library, top-performing 5′ UTRs contained more cap-proximal Us and fewer Gs than average, while the opposite was true of 5′UTRs with low activity (**Figure 3N**).

### Coding sequences significantly affect ribosome recruitment

The coding region of therapeutic mRNA is largely dictated by the desired protein product, with some room to optimize synonymous codons. In contrast, the 5′ UTR can be changed to suit the payload. Given that the scanning 48S complex contacts 6-24 nts of CDS^17^, we wondered whether 5′ UTRs are equally active when paired with different coding sequences. We tested 4,340 5′ UTRs in two contexts: upstream of the EGFP coding sequence or the endogenously occurring CDS for each 5′ UTR (**Figure 4A**). We noted that changing the coding sequence caused substantial differences in ribosome recruitment when comparing identical 5′ UTRs with EGFP or endogenous coding sequence (R^2^ = 0.48, **Figure 4B**). Among the 5′ UTRs tested with both coding sequences, nearly 20% (855) exhibited over a 2-fold change in ribosome recruitment activity (**Figure 4C**). We observed similar large effects on initiation upon replacement of EGFP with Fluc or SARS-CoV2 spike protein coding sequences (**Figure S4**). These data indicate that identifying the most optimal sequence is likely to require screening with the desired coding sequence.

**Figure 4.**
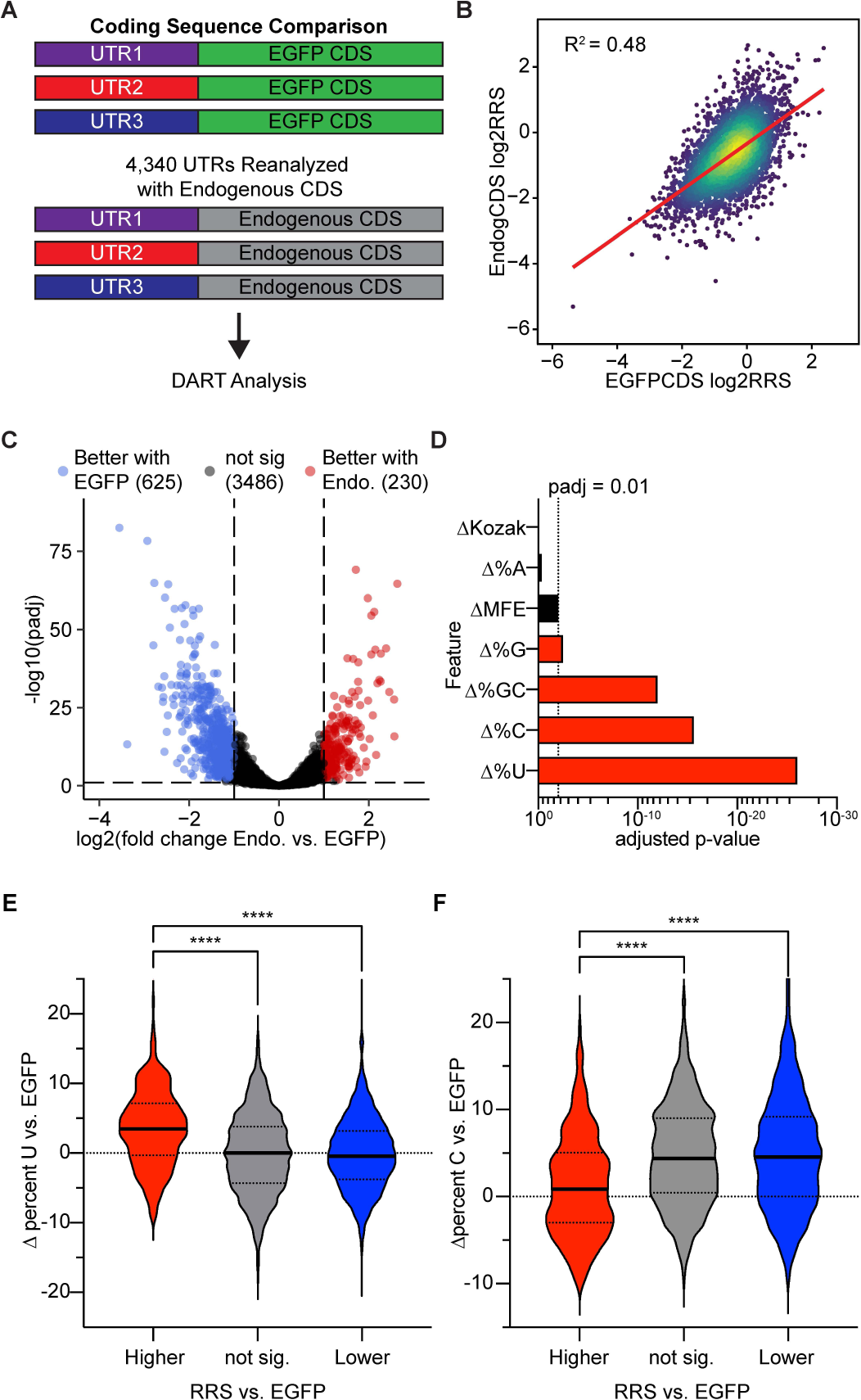
Coding sequence significantly affects ribosome recruitment to many 5′ UTRs. A) Library design to test the impact of coding sequences on ribosome recruitment. B) Coding sequence significantly affects ribosome recruitment. Each 5′ UTR is plotted according to its ribosome recruitment with EGFP and endogenous coding sequence. C) The impact of coding sequence on ribosome recruitment is context-dependent. 625 5′ UTRs exhibit significantly higher ribosome recruitment with EGFP coding sequence (blue), while 230 5′ UTRs have significantly higher ribosome recruitment when paired with the corresponding endogenous coding sequence (fold change > 2, Benjamini-Hochberg adjusted p < 0.01). D) Changes in uridine and cytidine content within the coding sequence correlate with altered ribosome recruitment. P-values determined by Wilcoxon rank test and adjusted using the Benjamini-Hochberg method. E and F) Uridines or cytosines in the coding sequence promote or repress recruitment, respectively. Sequences were binned based on their ribosome recruitment activity relative to the corresponding 5′ UTR with the EGFP coding sequence. ****p < 0.0001 unpaired Welch’s t-test.

We next analyzed the impact of CDS features on ribosome recruitment and found that nucleotide content within the 5′ region of the coding sequence was significantly correlated with the changes in translation activity (**Figure 4D**). Specifically, coding sequences that outperformed their EGFP counterparts contained more Us and fewer Cs (**Figures 4E and 4F**), suggesting that the repressive and activating effect of C- and U-rich elements, respectively, (**Figures 2 and 3**) is not restricted to the 5′ UTR. Engineering the 5′ end of coding sequences according to these principles may therefore be a mechanism to increase the protein production of therapeutic mRNAs beyond maximizing codon optimality (see Discussion).

### DART can be miniaturized and multiplexed for higher throughput with less input

Our results establish DART as a facile method for quantitative analysis of cis-regulatory elements in human 5′ UTRs. In principle, DART could dissect 5′ UTR-specific responses to trans factor manipulations including knockdowns, transfections, and small molecule treatments. However, the initial DART conditions used 250 µL of cell extract in 500 µL reactions and required a separate centrifuge bucket for each sample, limiting throughput. We therefore tested DART performance with decreasing input material ranging from 100 µL of cell extract (∼10 million cells) to 10 µL (∼1 million cells). Barcodes were added prior to pooling reactions for centrifugation. The smallest scale (10 µL) maintained 87-89% coverage across the 5′ UTR pool (38,191 sequences) (**Figure 5A**) and reproduced the results from larger reactions (**Figure 5B**). Ribosome recruitment scores from miniaturized DART reactions accurately predicted translation output from luciferase reporters (**Figure 5C**). Thus, DART can be miniaturized to work with limited starting material. Using smaller reaction volumes along with the barcoding strategy also allows pooling multiple samples onto a single sucrose gradient, easily increasing throughput by tenfold.

**Figure 5.**
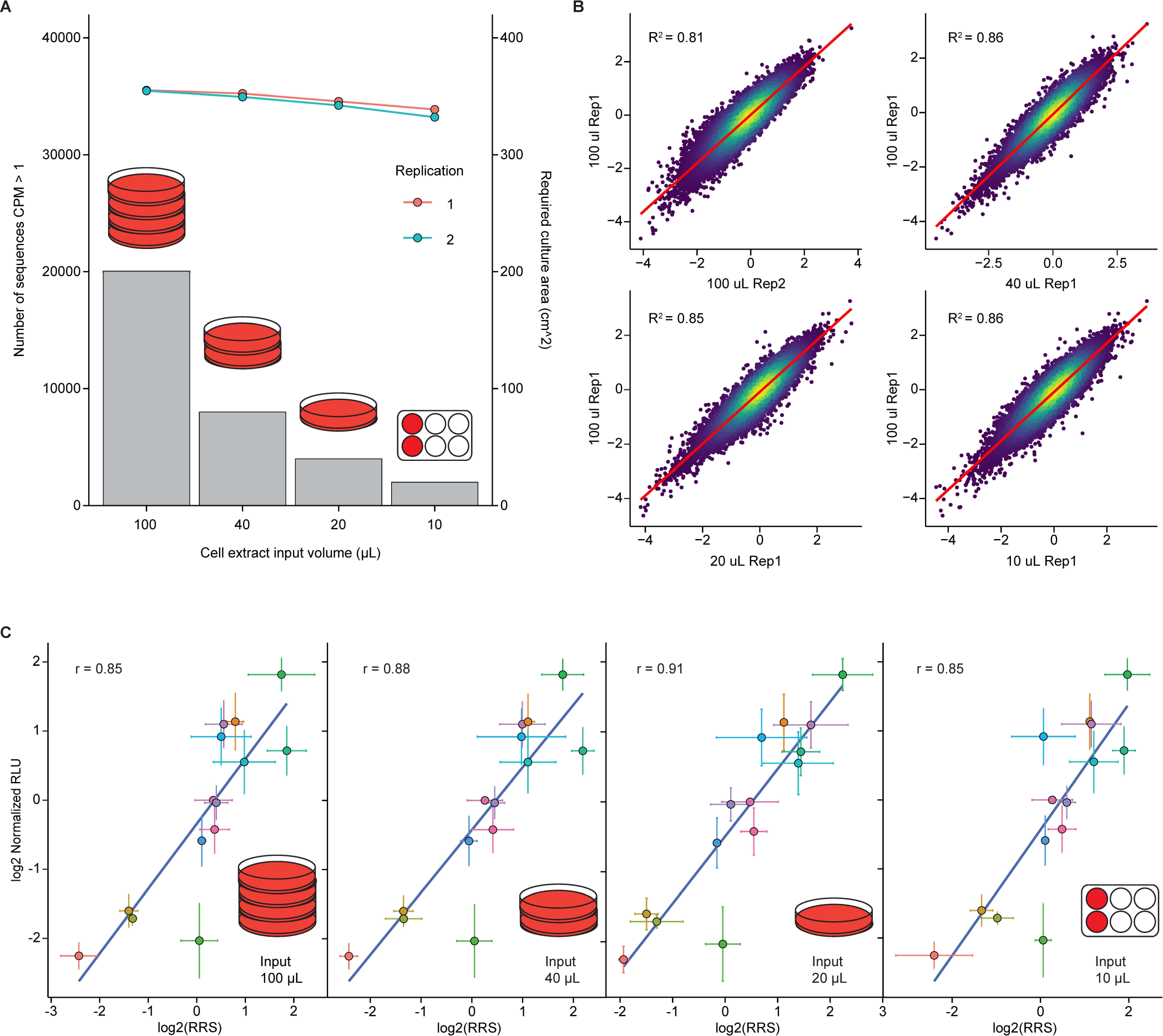
DART assay can be miniaturized to save input material and increase throughput. A) DART reaction can be scaled down 25-fold without loss of sequence representation. The amount of cell lysate input is plotted on the right y-axis with the number of sequences greater than 1 count per million plotted on the left y-axis. Cell culture plates indicate the amount of cell culture area required for each lysate input tested. B) DART reproducibility is preserved across miniaturization volumes. Ribosome recruitment scores measured from each input volume are plotted against each other. C) Ribosome recruitment scores obtained from miniaturized DART predict 5′ UTR-driven translational activity in full-length luciferase reporter mRNAs.

### N1-methylpseudouridine incorporation into 5′ UTRs significantly alters ribosome recruitment

Current-generation mRNA therapeutics are chemically substituted with N1-methylpseudouridine (m1Ψ) in place of uridine to avoid the innate immune response which would otherwise depress protein production *in vivo*^10,34^. Beyond this global effect, it was unclear how m1Ψ within 5′ UTRs would impact translation initiation. We tested this systematically by performing DART on 5′ UTR sequences with full replacement of uridine with m1Ψ. Importantly, uridine and m1Ψ-containing RNAs were co-incubated in the same translation reaction to allow the identification of sequence-dependent direct effects of m1Ψ substitution separate from any global effects (**Figure 6A**). In contrast to the negligible inter-replicate variability of ribosome recruitment activity by uridine- and m1Ψ-containing 5′ UTRs (R^2^ = 0.83 – 0.96), we observed large differences in RRS scores when comparing identical sequences with or without m1Ψ substitution (R^2^ = 0.69, **Figure 6B**). The impact of general features (length, secondary structure, GC content, Kozak strength, and the impact of cap-proximal nucleotides) was similar in m1Ψ-substituted 5′ UTRs (**Figure S5**). Surprisingly, we observed a global stimulatory effect of m1Ψ substitution on ribosome recruitment (1.9-fold on average), with over 36% of 5′ UTRs (n = 13,608) exhibiting over a 2-fold increase in initiation activity compared to unmodified sequences (**Figures 6C and 6D**). We confirmed that m1Ψ incorporation did not significantly alter the stability of the RNA pool during the translation reaction (**Figure 6E**), demonstrating that m1Ψ directly enhances ribosome recruitment.

**Figure 6.**
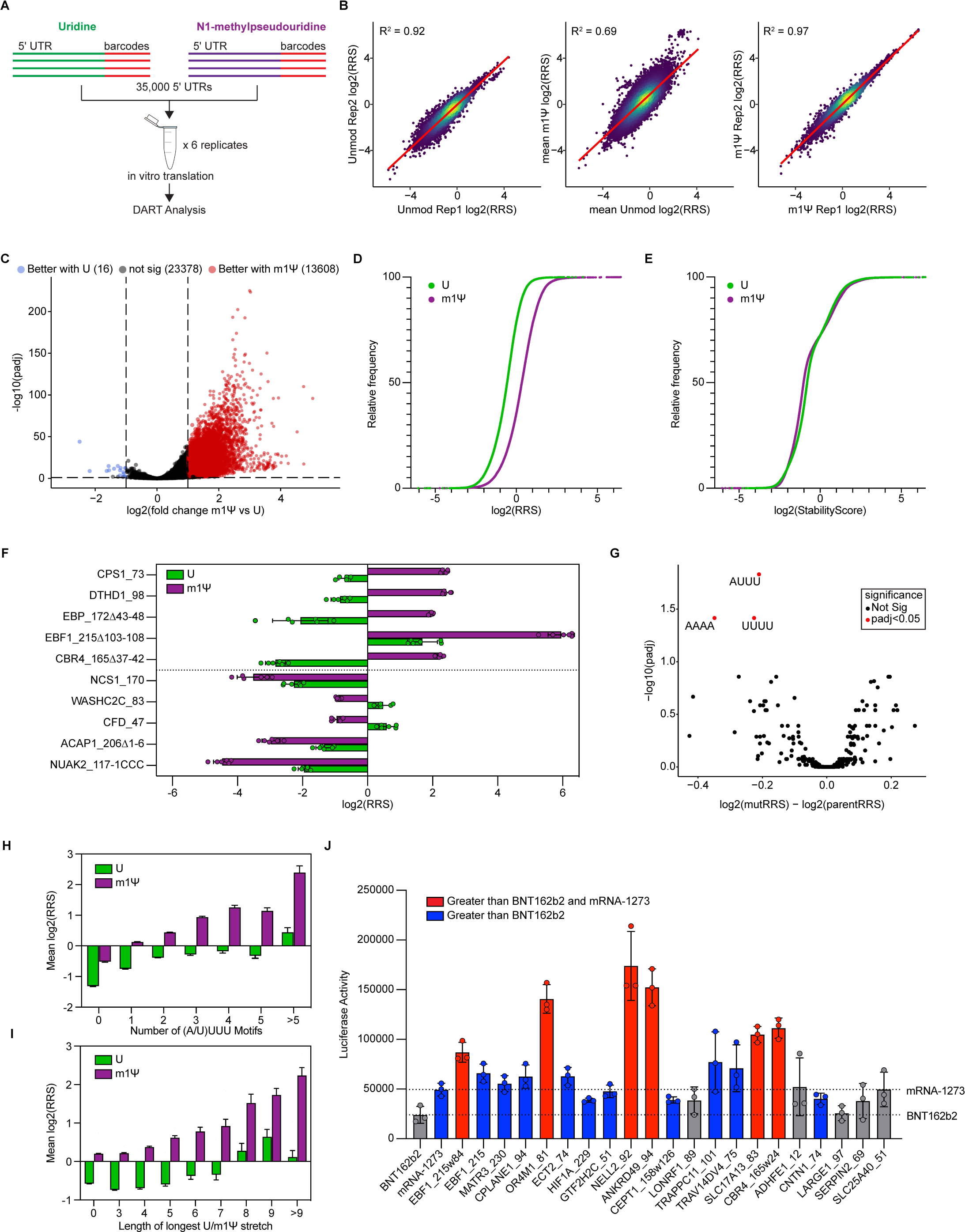
Global and 5′ UTR-specific effects of N1-methylpseudouridine on ribosome recruitment. A) Barcoding library design for testing the impact of N1-methylpseudouridine (m1Ψ) on 5′ UTR activity. Uridine-containing and m1Ψ-containing RNAs are barcoded and combined in the same reaction to allow direct comparison of activity. B) Identical 5′ UTR sequences containing uridine versus m1Ψ (middle) show widespread differences in ribosome recruitment (average of six replicates) compared to replicate reproducibility with uridine (U, left) or N1-methylpseudouridine (m1Ψ, right). C and D) m1Ψ globally promotes ribosome recruitment. C) Volcano plot of ribosome recruitment differences with m1Ψ substitution 13,608 5′ UTRs exhibit significantly reduced ribosome recruitment when uridine is replaced with m1Ψ (red), while 16 5′ UTRs show significantly increased ribosome recruitment with m1Ψ (blue, fold change > 2, Benjamini-Hochberg adjusted p < 0.01). D) Cumulative distribution function plot of ribosome recruitment scores from UTRs containing uridine (green) or m1Ψ (purple). E) RNA stability in DART is unaffected by m1Ψ substitution. Cumulative distribution plot of stability scores from UTRs containing uridine (green) or m1Ψ (purple). F) The effect size of m1Ψ on ribosome recruitment is 5′ UTR-specific. Individual examples of 5′ UTRs that are strongly stimulated (top) or repressed (bottom) by m1Ψ substitution (n = six replicates of DART-determined log2(RRS) values). G) Scanning deletion analysis identifies tetramers that significantly alter ribosome recruitment when deleted in unmodified m1Ψ-substituted RNAs (right). H) Increasing numbers of (A/U)UUU motifs correlate with increased ribosome recruitment in m1Ψ-substituted RNAs. 5′ UTRs were binned based on the number of (A/U)UUU motifs they contain. I) Longer stretches of poly(m1Ψ) within 5′ UTRs correlate with increased ribosome recruitment. 5′ UTRs were binned according to the longest poly(m1Ψ) stretch they contain. J) Optimal m1Ψ-substituted mRNAs produce more protein when placed into luciferase reporter mRNAs than current commercial vaccine 5′ UTRs. Blue, more luciferase produced compared to BNT162b2 5′ UTR; Red, more luciferase produced compared to BNT162b2 and mRNA-1273 5′ UTRs, p < 0.05 by two-tailed Student’s t-test.

Beyond the global effect, the degree to which m1Ψ incorporation altered ribosome recruitment was highly sequence-specific with individual sequences ranging from 32-fold stimulation to 5-fold repression (**Figure 6F**). To determine the sequence elements responsible for the effect of m1Ψ substitution on 5′ UTR activity, we performed DART analysis on the scanning deletion library (**Figure 2A**) with and without m1Ψ substitution. This analysis revealed that deletion of (A/X)XXX (X=m1Ψ) motifs significantly reduced ribosome recruitment activity (**Figure 6G**). Notably, deletion of these motifs did not have a significant impact on unmodified 5′ UTRs (**Figure 2D**), indicating these sequence elements are stimulatory only when modified to m1Ψ. Accordingly, across all sequences tested with m1Ψ, increasing numbers of (A/X)XXX motifs correlated with increased ribosome recruitment (**Figure 6H**). As the (A/X)XXX elements contain consecutive m1Ψ nucleotides, we were curious whether longer stretches of poly(m1Ψ) affected ribosome recruitment. We noted a significant increase in ribosome recruitment to 5′ UTRs containing longer stretches of poly(m1Ψ) (**Figure 6I**). In contrast, the length of unmodified poly(U) stretches had little effect.

Finally, we sought to validate candidate 5′ UTRs that could drive higher levels of protein synthesis than current mRNA vaccines. We selected 21 of the top-performing 5′ UTRs from the m1Ψ DART screen, cloned them in front of luciferase reporters, and transfected these reporter mRNAs into Hela cells. 16/21 and 6/21 outperformed the 5′ UTR sequences from the FDA-approved SARS-Cov2 vaccines BNT162b2 (Pfizer-BioNTech) and mRNA-1273 (Moderna), respectively (**Figure 6J**), demonstrating the utility of the DART method to identify highly active sequences for mRNA therapeutic design.

## Discussion

Here, we present the development of DART as a high-throughput method to quantify the translational activity of 5′ UTRs in a human system. A major advantage of the DART assay is the ease with which 5′ UTR libraries can be designed and analyzed. Regulatory features identified in an initial screen can be subsequently interrogated directly, moving from correlation to causation. We use this approach to dissect intrinsic determinants of large differences in translation initiation efficiency, quantifying the impact of known regulatory features while identifying novel design principles for therapeutic mRNA development.

Optimization of therapeutic mRNAs typically includes uridine depletion and modification of the remaining uridines to N1-methylpseudouridine—both aimed at reducing activation of the cellular innate immune response^34,35^. Our results demonstrate a more nuanced approach is necessary. By isolating the initiation step of translation, we find that uridines within the cap-proximal region of 5′ UTRs significantly enhance ribosome recruitment. This effect holds when uridines are substituted with m1Ψ. Surprisingly, however, the substitution of uridine with m1Ψ globally increases ribosome recruitment, indicating unique effects of m1Ψ compared to uridine. We observe that stretches of consecutive m1Ψ—but not uridine—residues within 5′ UTRs have a substantial enhancing effect on translation activity. Employing a sweeping uridine depletion strategy is therefore likely to remove potent translational enhancers and decrease protein output from therapeutic mRNAs.

In addition to uridine content optimization, our data reveals that cytosine-rich elements can function as potent translational silencers. Notably, CCC elements repress ribosome recruitment while also inhibiting enzymatic mRNA capping by base-pairing with 5′ end and rendering it inaccessible to the Vaccinia capping enzyme. Some current-generation mRNA therapeutics, including mRNA-1273, use an enzymatic capping strategy^36^. Our results indicate that sequence choice can impact mRNA production as well as translational activity in cells. Interestingly, we find that the enhancing and repressive effects of uridine and cytosine, respectively, observed in the 5′ UTR also extend to the coding sequence. This may provide another mechanism to optimize the coding sequence in addition to codon optimality.

Beyond therapeutic mRNA development, DART has broad applications to examine effects on translation initiation in other contexts. Multiple small molecules in pre-clinical or clinical-stage development for cancer treatment, including silvestrol and zotatafin (eIF4A inhibitors), ribavirin and 4Ei-1 (eIF4E inhibitors), and everolimus and BEZ235 (mTOR inhibitors) are designed to inhibit or alter translation initiation^37,38^. The ability to perform DART with minimal input material and multiplex multiple samples on a single sucrose gradient makes it straightforward to perform dose-escalation studies to define drug-sensitive and -insensitive 5′ UTRs. Moreover, cancerous tissues reprogram their transcriptome, producing an abundance of tumor-specific 5′ UTR isoforms^39,40^. Our results demonstrate that relying on simple principles such as 5′ UTR length, structure formation, etc. is insufficient to predict the translational impact of these isoform changes. DART is a facile method to empirically measure the consequences of UTR variation and test the impact of therapeutic agents on the disease-specific transcriptome.

## Limitations

Here, we present DART as a powerful approach to study translation initiation by human 5′ UTRs. The method does possess certain limitations. First, DART requires translationally active extract from the cell type or tissue of interest. We have successfully produced active extracts from a wide range of cancerous and immortalized mammalian cell lines. However, producing extracts from less translationally active inputs such as primary cells or tissue samples is likely to require optimization. Second, in vitro transcription with T7 and co-transcriptional capping with CleanCap AG yields RNAs that begin with 5′-AG-3′ immediately following the methylguanosine cap. Evaluating the impact of cap-proximal nucleotide positions would require changes to the method of synthesizing the RNA pool. Third, in this study, we fully replaced uridine with m1Ψ to match the design of therapeutic mRNAs. In future studies, site-specific modification can be achieved by incubating the pool RNA with recombinant modifying enzymes to uncover the function of endogenous 5′ UTR modifications.

## Materials and Methods

### 5′ UTR Pool Design and Synthesis

To obtain the 5′ UTRs that are expressed in human tissue, total RNAseq datasets for liver, kidney, neuron, T cell, K562, MCF10a, and HepG2 from the ENCODE project were gathered and aggregated. Transcripts were first filtered by length ranging from 10-230 nucleotides and by TPM > 1 in any dataset. The top 32,355 transcripts with the highest TPM were selected. The top 6,000 transcripts were also chosen for the list of endogenous CDS. The sequences of the selected 5′ UTR and CDS were retrieved from Gencode41. The 5′ UTR sequences were concatenated with EGFP CDS sequences or their endogenous CDS sequences. T7 promoter sequence (GCTAATACGACTCACTATAGGG) was added in front of 5′ UTRs and RT primer binding sequence (CACTCGGGCACCAAGGAC) was added to the 3′ end. CDS sequences were trimmed from 3′ end to make 300-nucleotide total length oligos. All upstream ATGs in the 5′ UTRs were mutated to AGT. The last in-frame codon in CDS was changed to termination codon TAA.

The scanning deletion pool was constructed based on the top 200 and bottom 200 5′ UTR sequences from the initial DART results. For each parental sequence, a scanning deletion of every 6-nt window was made throughout the entire 5′ UTR to generate deletion variants. Upstream ATG generated from the deletion were removed by mutation to AGT.

Designed oligos were purchased as a DNA pool (Twist Bioscience) and PCR amplified with oXX01 (enzymatic capping) or oXX02 (co-transcriptional capping) and the PCR product was gel-purified. Pool RNA was produced by runoff T7 transcription using the MEGAshortscript T7 transcription kit (ThermoFisher AM1354) using the purified DNA template. The pool RNA was then gel-purified and capped using the Vaccinia Capping System (NEB M2080S). For co-transcriptional capping, the T7 promoter sequence was changed to GCTAATACGACTCACTATAAGG and CleanCap AG (TriLink N-7113) was added to the in vitro transcription reaction according to manufacturer protocols.

For multiplexed DART reactions, DNA pools were PCR amplified with the common oXX02 forward primer and barcoded reverse primers oXX_Bar1-8. Four barcode sequences were used for uridine and four were used for N1-methylpseudouridine modified RNAs. N1-methylpseudouridine substituted RNAs were produced by the complete replacement of uridine triphosphate in the T7 transcription reaction with N1-methylpseudouridine triphosphate (Trilink N-1081). Uridine and N1-methylpseudouridine RNAs were then mixed in equimolar ratios prior to in vitro translation.

### In vitro translation

For standard DART reactions, 40 picomoles of capped pool RNA was added to each 0.5 mL in vitro translation reaction containing 0.25 mL of Hela cytoplasmic extract (Ipracell CC-01-40-50), 16 mM HEPES KOH pH 7.4, 40 mM potassium glutamate, 2 mM magnesium glutamate, 0.8 mM ATP, 0.1 mM GTP, 20 mM creatine phosphate, 0.1 mM spermidine, 165 µg creatine phosphokinase, 1.6 mM DTT, 2mM PMSF, 140 units of RNasin Plus, 1X cOmplete EDTA-free protease inhibitor cocktail, and 0.5 mg/mL cycloheximide. Reactions were incubated at 37 °C for 30 minutes in a shaking thermomixer. Following incubation, reactions were immediately loaded onto a sucrose gradient for 80S isolation. Miniaturized DART reactions maintained the same concentrations of components.

### 80S Isolation and RNA extraction

Translation reactions were loaded onto 10-50% sucrose gradients containing 20 mM HEPES KOH pH 7.4, 2 mM magnesium glutamate, 100 mM potassium glutamate, 0.1% Triton X-100, 3 mM DTT, 0.1 mg/mL cycloheximide and centrifuged at 35,000 RPM for 3 hours at 4°C in a Beckman SW41 rotor. Gradients were fractionated using a Biocomp Gradient Station (Biocomp Instruments) with continual monitoring of absorbance at 260 nm. Fractions corresponding to the 80S peak were collected and pooled. To extract RNA from the pooled fractions, 650 µL of acid phenol and 40 µL of 20% SDS was added per 600 µL of fraction volume. The mixture was then transferred to a 65 °C water bath and incubated for 10 minutes, vortexing every minute. The samples were then cooled for 5 minutes on ice and transferred to a pre-spun 15 mL MaXtract tube (Qiagen 129065) containing an equal volume of chloroform to the acid phenol used. After centrifugation to separate the aqueous and organic phases, the aqueous phase containing RNA was transferred to a new 1.5 mL tube and subjected to a second extraction using 650 µL of phenol:chloroform:IAA (25:24:1) pH 6.8 per 600 µL of aqueous volume. A final extraction was performed using 650 µL of chloroform per 600 µL of aqueous volume and the extracted RNA was precipitated with isopropanol.

### DART Library Preparation

RNA from the input pool and 80S fractions was reverse transcribed using XXX primer using Superscript III (Invitrogen 18080093). The resulting full-length cDNA was gel-purified and ligated to the 5′ adaptor OWG920 with High Concentration T4 RNA Ligase (NEB M0437M) overnight on a shaking thermomixer. The ligated cDNA was then purified using 10 µL of MyOne Silane beads (Invitrogen 37002D). Libraries were then amplified using the primers XX and XX, gel purified, and sequenced on a HiSeq 2500.

### Data Processing

The raw fastq files were first demultiplexed using Flexbar^41^. After demultiplexing, each file was processed with BBMap module to trim adaptors, collapse duplications, and remove UMIs. The processed sequencing results were aligned to pool sequences using STAR^42^, with the following parameters: soloStrand = Forward; alignSJoverhangMin = 999; alignIntronMax = 1; alignIntronMin = 999; outFilterMismatchNmax = 1. The counting of alignment was performed by BBMap pileup. The sequences with < 10 counts in any of the samples were removed. To calculate counts per million (CPM), a total depth for each translation reaction was obtained by summing total counts from all the samples that derived from the same translation reaction, then the raw counts for each sample were normalized by this total depth.

### In vitro luciferase assays

5′ UTR sequences were cloned into a plasmid downstream of a T7 promotor and immediately upstream of GFP coding sequence followed by a 15 amino acid glycine-serine linker and either Nano luciferase or Firefly luciferase coding sequence and an encoded 60 nucleotide poly(A) tail. The resulting plasmids were then linearized and used as a template for run-off T7 transcription using the MEGAscript T7 transcription kit (Invitrogen AMB13345). 0.05 pmol of mRNA was incubated in a 10 µL in vitro translation reaction containing 4 µL of Hela cytoplasmic extract, 16mM HEPES KOH pH 7.4, 0.1 mM spermidine, 0.8 mM ATP, 0.1 mM GTP, 40 mM potassium glutamate, 2 mM magnesium glutamate, and 20 mM creatine phosphate. Reactions were incubated at 37 °C for 30 minutes in a shaking thermomixer. Following incubation, reactions were immediately halted with ice-cold Passive Lysis Buffer (Promega E1941). Firefly luciferase and nano luciferase levels were then measured using Bright-Glo (Promega E2620) and Nano-Glo (Promega N1130), respectively, on a luminometer.

### In cell luciferase assays

Hela cells were cultured in DMEM supplemented with 10% FBS. 10,000 cells per well were seeded into a 96-well plate and allowed to adhere overnight. The following day, cells were transfected with 100 nanograms of mRNA using the Lipofectamine MessengerMAX Transfection Reagent (Invitrogen LMRNA015). Cells were lysed in 50µL of Bright-Glo plus 50 µL of 1X phosphate-buffered saline and luciferase activity was measured on a luminometer.

### Tetramer analysis from scanning mutagenesis

The change in ribosome recruitment score (deltaRRS) was obtained by dividing the RRS for each deletion sequence by the RRS for its parent sequences. Tetramers contained within each 6-nucleotide list were associated with a list of all possible tetramer sequences. To reduce noise, only tetramers that were deleted at least 30 times were considered. The impact of each tetramer sequence was calculated as the mean of deltaRRS values. Significance was determined by Wilcoxon test and the p-values were adjusted using the Benjamini-Hochberg correction. P-values less than 0.01 were considered significant.

### DMS-MaPseq library preparation

DNA pool was amplified with PCR (forward primer oXX02; reverse primer oXX_Bar2. RNA was produced by T7 in vitro translation using the MEGAshortscript kit (ThermoFisher Scientific AM1354), co-transcriptionally capped with CleanCap Reagent AG (TriLink N-7113), treated with TURBO DNase (Invitrogen) and gel-purified. For DMS probing, RNA (2 μg in 6 μL water) was denatured at 95 °C for 2 min. RNA was refolded by adding 88.8 μL 300 mM sodium cacodylate and 1 μL RNasin Plus (Promega N2611) and 10 min incubation at 37 °C, followed by addition of 1.2 μL 500 mM MgCl2 and 20 min incubation at 37 °C. DMS (3 μL, Sigma-Aldrich D186309) was added to the folded RNA solution and allowed for 2 min incubation at 37 °C, then quenched by adding 42.8 μL β-mercaptoethanol (BME). For denatured RNA control, RNA (2 μg in 6 μL water) was mixed with 1 μL RNasin Plus, 39.2 μL water, 50 μL formamide (Invitrogen 15515-026) and 0.8 μL 0.5 M EDTA and incubated at 95 °C for 2 min. The solution was incubated with 3 μL DMS and quenched with 42.8 μL BME. RNAs were purified via ethanol precipitation and reverse-transcribed using TGIRT-III (InGex) and primer oXX04. The resulting cDNA was gel purified and a 5’ adaptor (oXX03) was ligated. Libraries were prepared as described above and sequenced via Novaseq.

### DMS-MaPseq analysis

The data processing steps for DMS-MaPseq results, including adapter removal, PCR duplicate removal, and alignment follow the procedure for DART analysis, except using the default setting for STAR: outFilterMismatchNmax. Mismatches to the reference sequence were labeled with MD tags using the SAMtools calmd function. The coverage and mutation rate for each position were calculated and averaged within replicates using a house-developed script. The sequences with a coverage of < 200 reads were removed from further analysis. For each sequence, the mutation rates on A/C positions were normalized to a 0-1 scale, while U/G positions were not considered. The mutation rates were used to predict base pairing probability using RNAfold from the ViennaRNA package.

### Linear regression model

The linear regression model was built with the R caret package based on four features: 5′ UTR length, GC content in 5′ UTR, the minimum free energy, and the Kozak similarity score. DART results were randomly split, 80% (29523 5′ UTRs) for the training set and 20% (7396 5′ UTRs) for the test set. The accuracy of the linear model was determined by the correlation between the prediction and the RRS scores observed in the test set.

### Preparation of HEK293T translation extracts

HEK293T cells were maintained in DMEM (Gibco 11965-092) supplemented with 10% FBS (heat-inactivated, Sigma F4135). The cells were seeded at 17,200 cells/cm^2^ and cultured for 3 days. At the moment of harvest, the cells were dissociated with 0.25% trypsin-EDTA (Gibco 2520056), and the collected cell pellet was washed once with 30 mL of ice-cold PBS and once with 10 ml ice-cold isotonic buffer (HEPES-KOH 16 mM, potassium acetate 100 mM, magnesium acetate 0.5 mM, DTT 5 mM). The pellet was resuspended and lysed with an equal volume of hypotonic buffer (HEPES-KOH 16 mM, potassium acetate 10 mM, magnesium acetate 0.5 mM, DTT 5 mM), and incubated on ice for 10 min. The cell lysate was homogenized with 10 passes through a 27-gauge needle and centrifuged at 16,000 x g at 4°C for 1 min. The supernatant was collected and frozen at −80°C until use.

### Analysis of differential ribosome recruitment

Differential ribosome recruitment activity between 5′ UTRs paired with their endogenous coding sequence and their EGFP counterparts or between unmodified and m1Ψ-substituted RNAs was determined using the R package DESeq2^43^. For comparisons between U and m1Ψ, the read depths for each sample were used to estimate size factors. P-values were adjusted using Benjamini-Hochberg correction and an adjusted p-value of less than 0.01 and an absolute log2 fold change > 1 were considered significant changes.

## Acknowledgments

We thank all members of the Gilbert and Thoreen labs for discussions and critical reading of the manuscript. This work was supported by the U.S. National Institutes of Health (R01GM132358 to W.V.G., F31DK129022 to C.J.L., F31CA254339 to A.S.D.), the American Cancer Society (PF-23-1144579-01-RMC to D.J.), the National Science Foundation (G.R.F.P. to K.S.A.) and a Sponsored Research Agreement with Pfizer (ITEN2021.RNA01 to W.V.G. and C.C.T.). The funders have no role in the design or conduct of the study; collection, management, analysis, or interpretation of the data; preparation, review, or approval of the manuscript; or the decision to submit the manuscript for publication.

## Author Contributions

C.J.L. and W.V.G. conceived the project. C.J.L., L.X., S.B., D.J., and K.S.A., performed experiments. C.J.L., L.X., S.B., D.J., A.S.D., Y.C., and C.C.T. analyzed high-throughput sequencing data. W.V.G. and C.C.T. supervised the study. C.J.L. wrote the first draft of the manuscript. All authors discussed the results and edited the manuscript.

## Ethics Declarations

Yale University has filed a patent application on the basis of this work. C.J.L., L.X., C.C.T., and W.V.G., are named as co-inventors.

## Data Availability

Raw and analyzed data from all high-throughput sequencing experiments have been deposited on the Gene Expression Omnibus (https://www.ncbi.nlm.nih.gov/geo/) under accession number GSE256185.

## Code Availability

All scripts used in the analysis of DART data are available at https://github.com/carsonthoreen/dart_2024

---

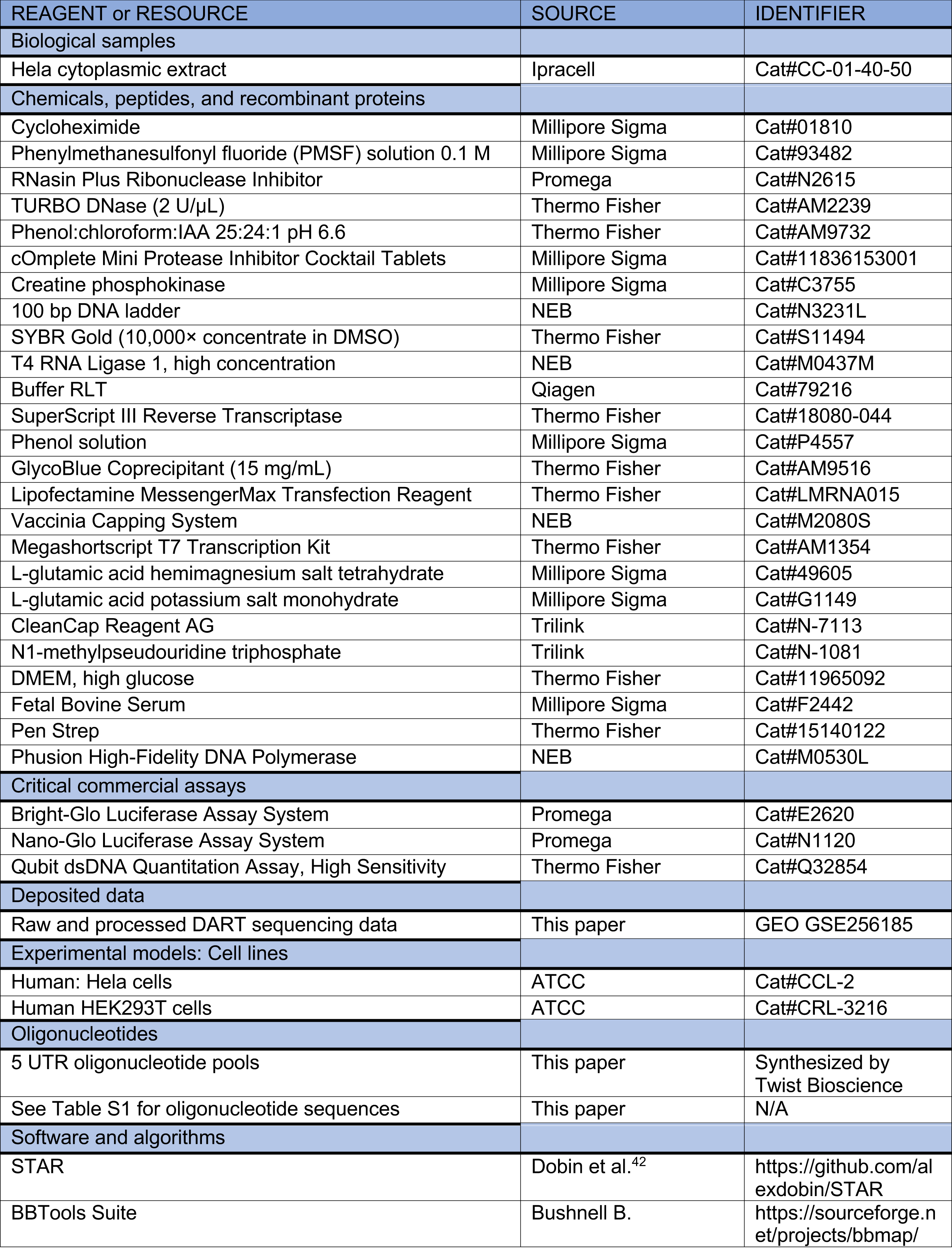
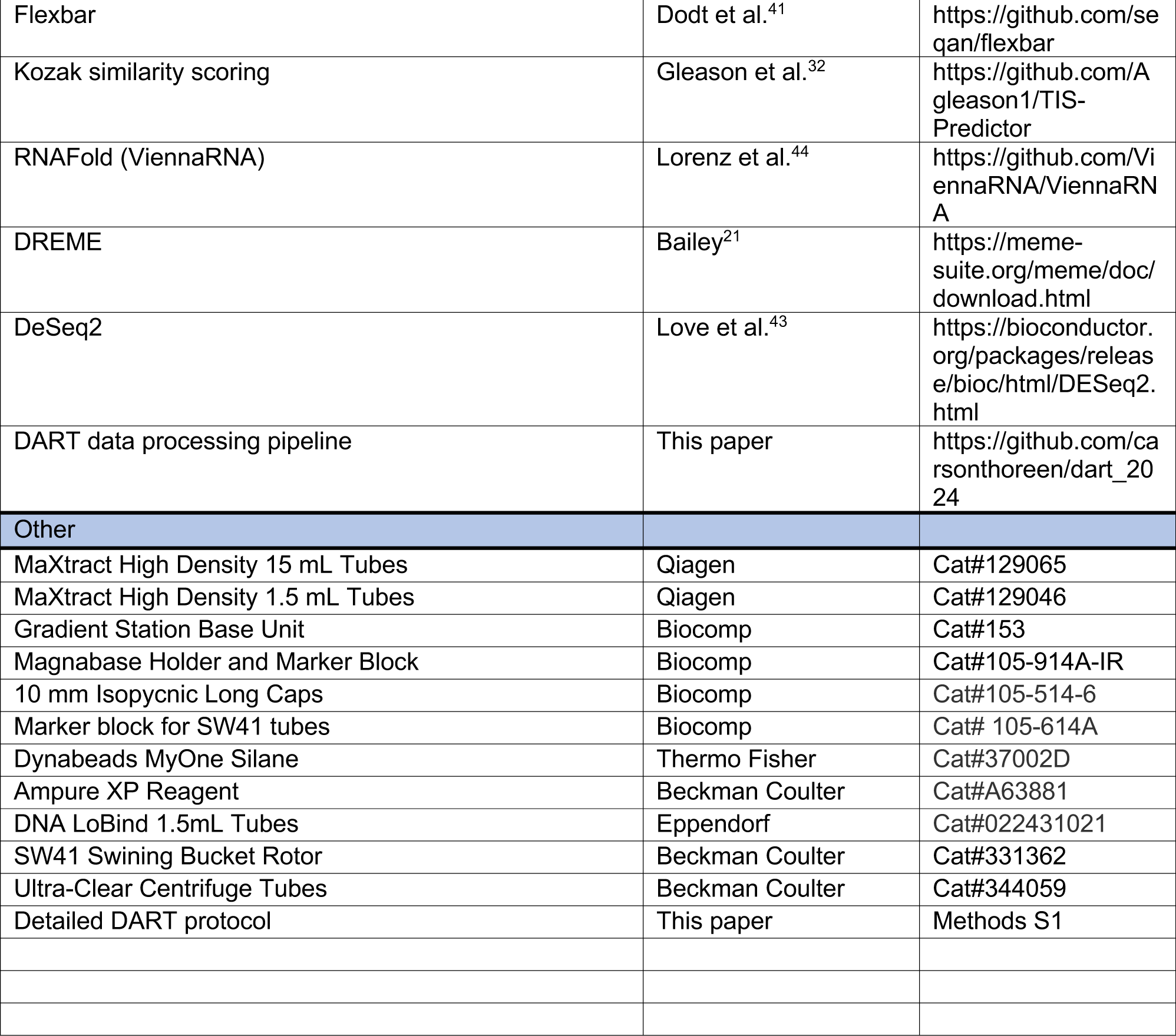

**Table S1. Oligonucleo1des used for DART library synthesis and sequencing**

**Table S2. RNA sequences, counts per million, and ribosome recruitment scores for Vaccinia-capped RNAs**

**Table S3. RNA sequences, counts per million, and ribosome recruitment scores for co-transcrip1onally-capped RNAs**

**Figure S1.**
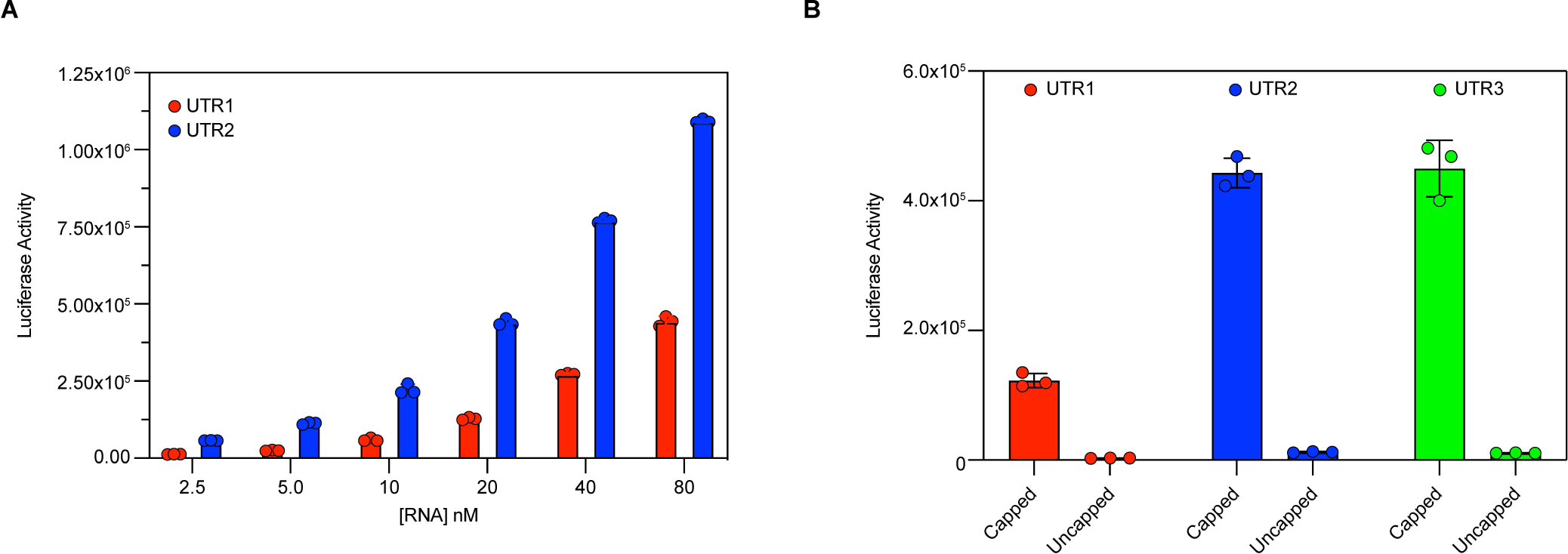
HeLa extract is active over a wide concentration of RNA and cap-stimulated. A) Luciferase production from reporter mRNAs scales with mRNA concentration from 2.5 to 80 nanomolar. B) Addition of an N7-methylguanosine cap increases protein production from three different luciferase mRNAs (UTR1, 41.4-fold; UTR2, 36.2-fold; UTR3, 41.9-fold increase plus cap).

**Figure S2.**
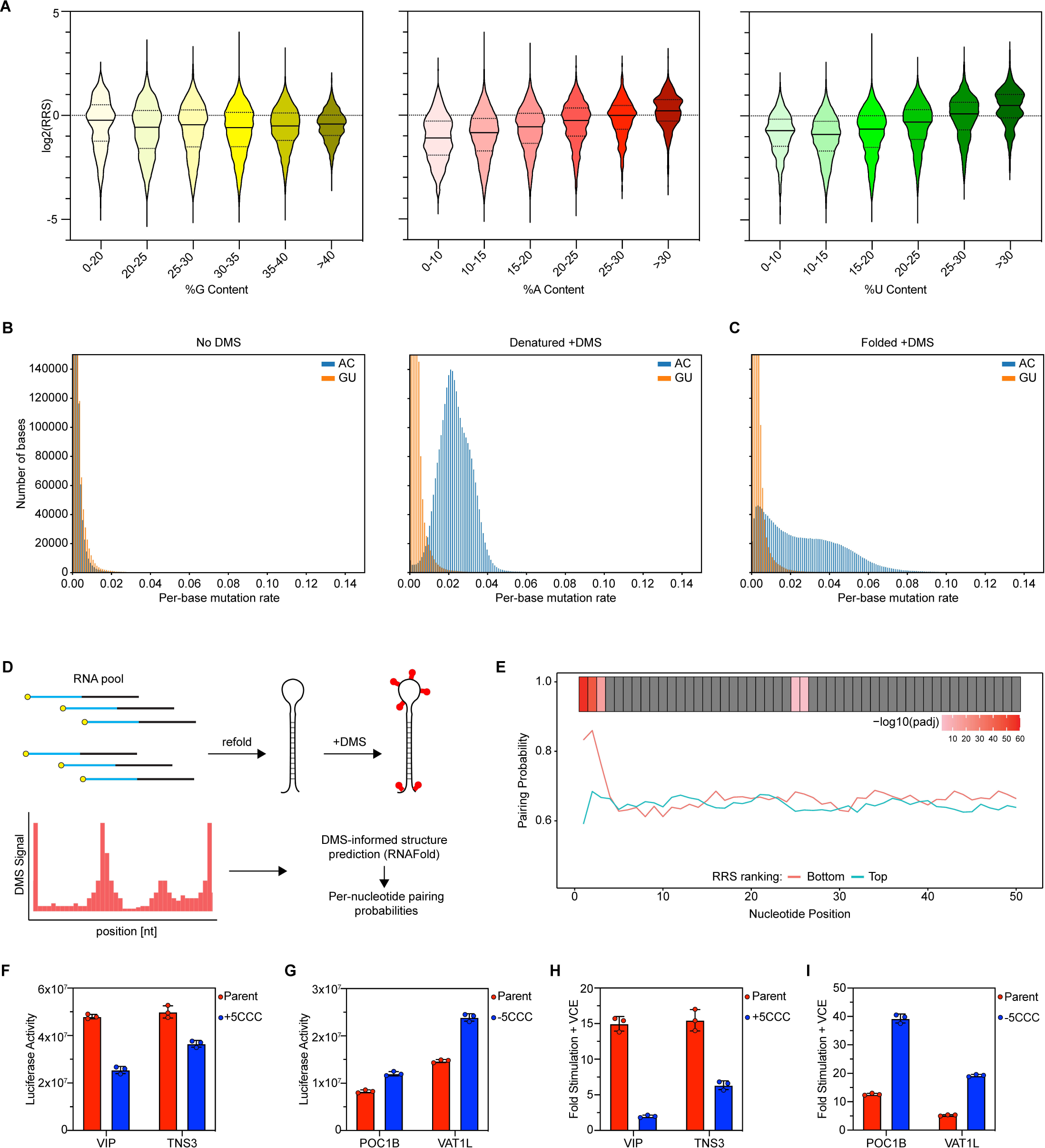
Effects of nucleotide content, RNA structure, and CCC motifs on translation. A) Global trends of ribosome recruitment with increasing guanosine (left), adenosine (middle), and uridine (left). 5′ UTRs binned by percent nucleotide content. B) DMS treatment specifically induces mutations at A and C in denatured RNA (right) compared to untreated control (left). C) Refolding RNA reduces DMS reactivity. D) DMS processing methodology for computing nucleotide pairing probabilities. E) Comparison of DMS-informed pairing probabilities for the top 10% (cyan) and bottom 10% (red) of 5′ UTRs by log2(RRS). Benjamini-Hochberg adjusted p-values shown (top) for each nucleotide. F) Effects of CCC motif addition and G) deletion on reporter RNAs capped with CleanCap AG. H) and I) Fold increase in luciferase production from reporter mRNAs capped with Vaccinia capping enzyme (VCE). The fold stimulation is shown for H) CCC motif addition and I) CCC motif deletion.

**Figure S3.**
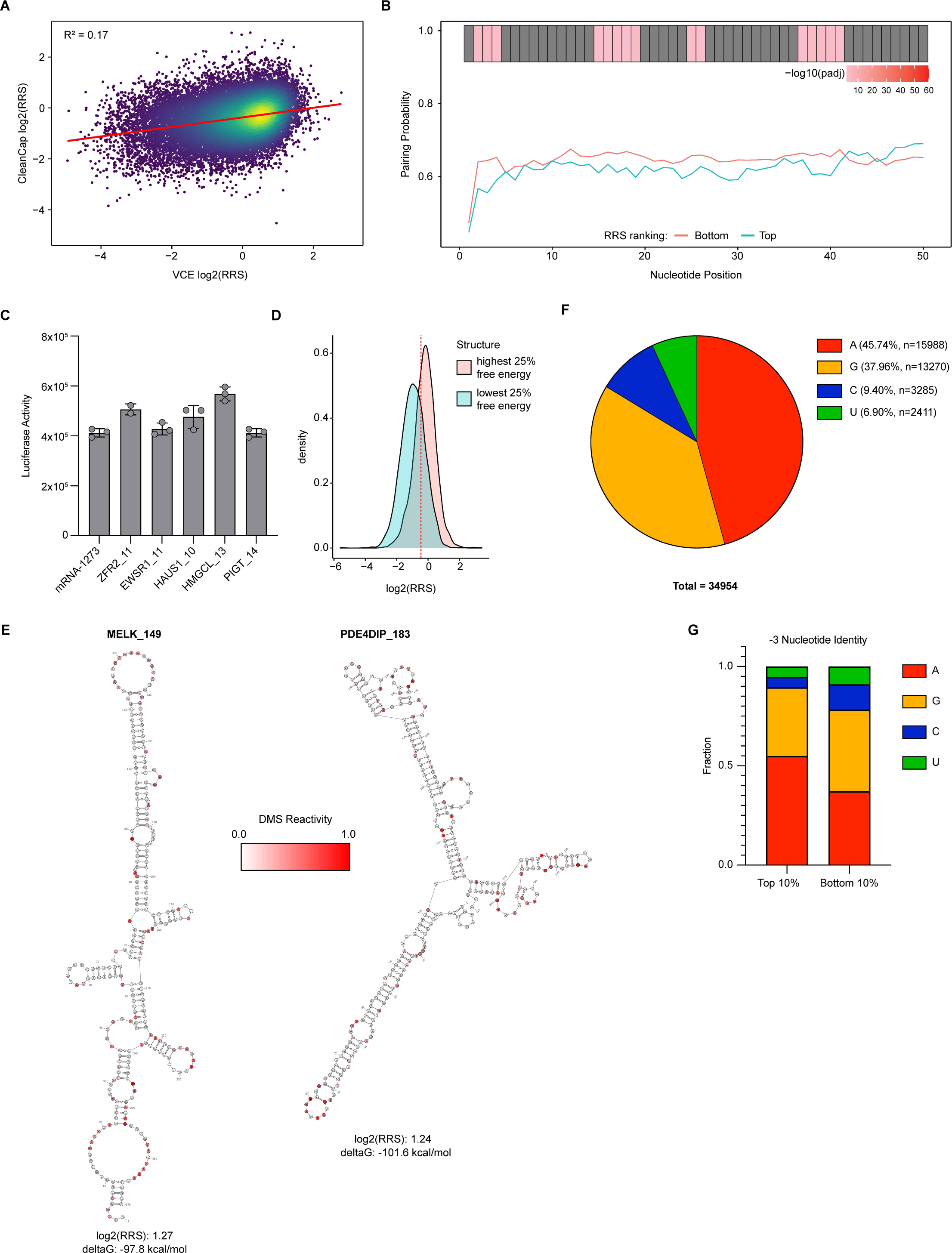
Regulatory features of 5′ co-transcriptionally capped 5′ UTRs. A) Comparison of ribosome recruitment scores from 5′ UTRs capped with Vaccinia capping enzyme (VCE) or co-transcriptionally capped with CleanCap AG. B) Comparison of DMS-informed pairing probabilities for the top 10% (cyan) and bottom 10% (red) of 5′ UTRs by log2(RRS). Benjamini-Hochberg adjusted p-values shown (top) for each nucleotide. C) Highly active short 5′ UTRs by DART analysis produce high levels of luciferase from reporter mRNAs. D) Histogram of log2(RRS) by 5′ UTRs binned by the lowest (cyan) or highest (red) quartile of minimum free energy. Dashed line indicates the median log2(RRS) from all 5′ UTRs tested. E) DMS-MaPseq informed RNA folding predictions of highly structured 5′ UTRs that were highly active in DART by log2(RRS). Nucleotides are shaded according to their reactivity in DMS MaPseq. F) Nucleotide identity of the −3 position relative to the start codon from all 5′ UTRs tested in DART. Nucleotide identity of the top 10% and bottom 10% of 5′ UTRs by log2(RRS) at the −3 position.

**Figure S4.**
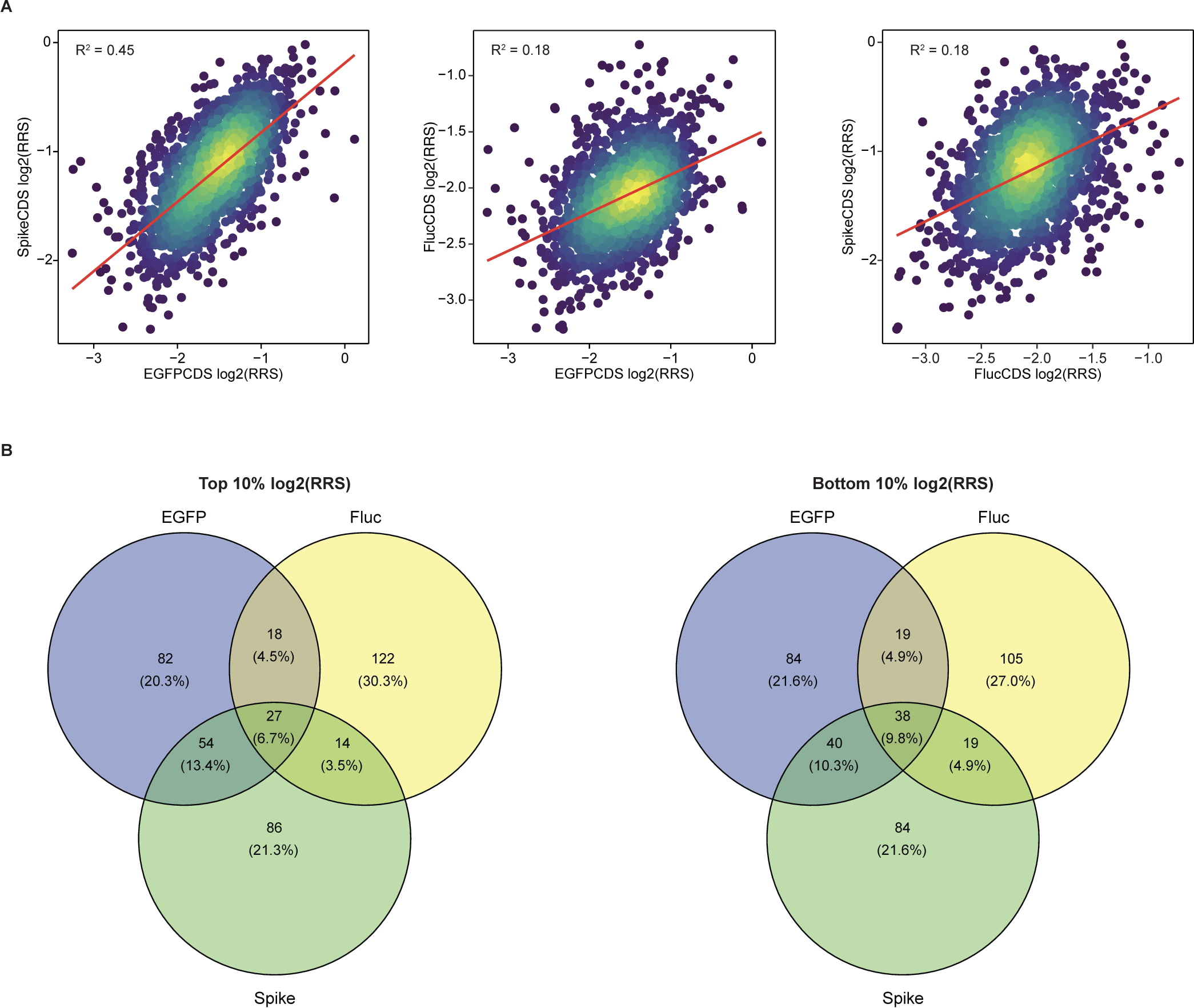
Coding sequences alter initiation efficiencies. A) Coding sequence significantly affects ribosome recruitment. Each 5′ UTR is plotted according to its ribosome recruitment with the indicated coding sequence (Spike: SARS-CoV2 spike protein; Fluc: firefly luciferase). Each coding sequence tested with 2,000 random 10-nucleotide 5′ UTRs. B) Minimal overlap between most (left) and least (right) active 5′ UTRs with different coding sequences. 5′ UTRs were binned into top and bottom deciles according to their ribosome recruitment scores measured by DART.

**Figure S5.**
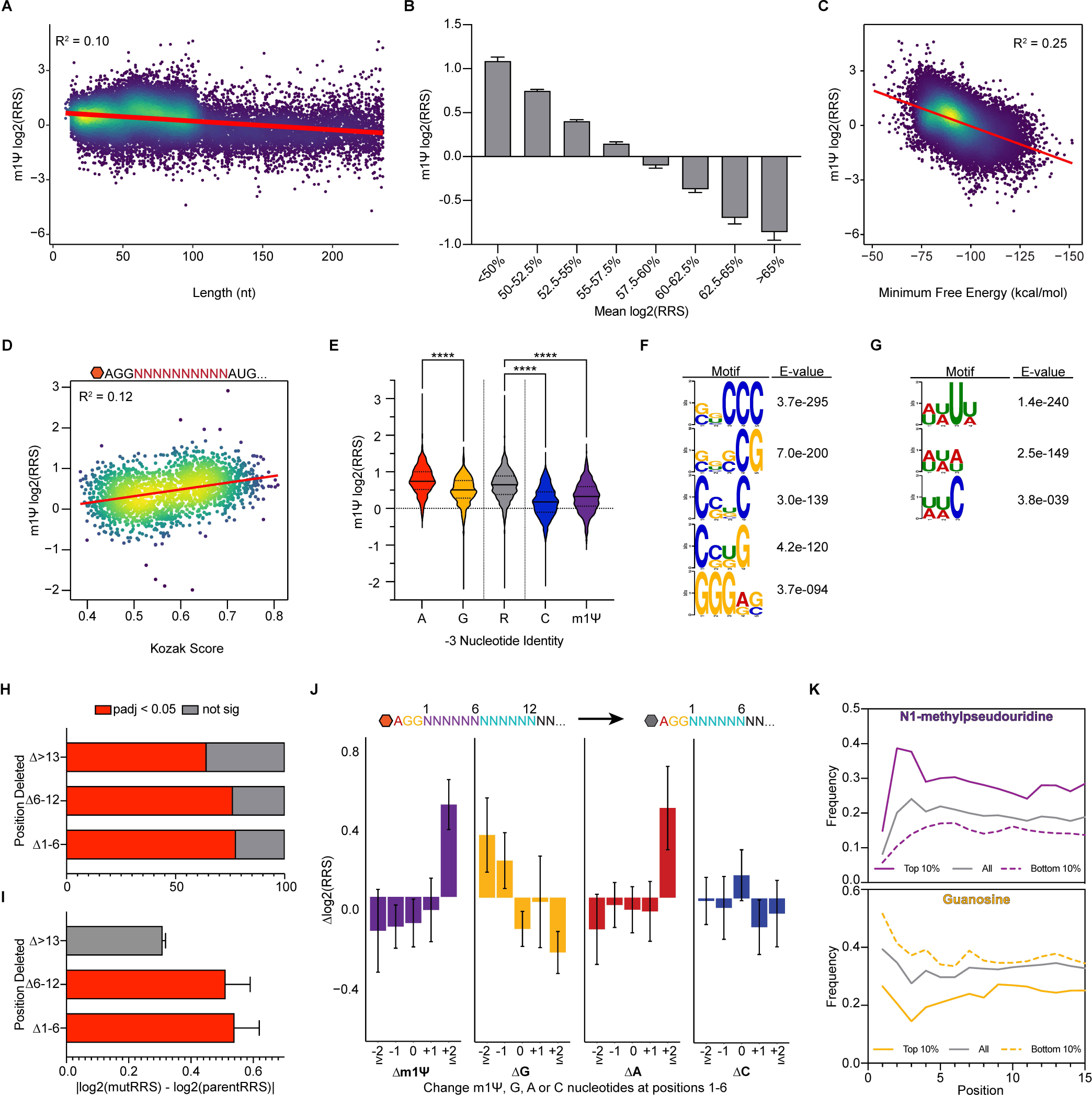
Impact of features on m1Ψ-substituted 5′ UTR activity. A) Effect of UTR length on ribosome recruitment. 5′ UTRs binned according to length. Data are represented as mean log2(RRS) for each bin (error bars = 95% CI). B) Impact of GC content on ribosome recruitment. Sequences were binned according to their percent GC. Data are represented as mean log2(RRS) for each bin (error bars = 95% CI). C) Predicted minimum free energy negatively correlates with ribosome recruitment and explains 25% of the variability observed in DART. D) Kozak sequence strength promotes ribosome recruitment to random 10-nucleotide 5′ UTRs. UTRs scored based on conformity to the consensus human Kozak sequence and plotted against log2(RRS). E) Pyrimidines at the −3 position correlate with worse ribosome recruitment. 5′ UTRs were binned based on their nucleotide identity at −3 relative to the start codon. ****p < 0.0001, unpaired Welch’s t-test. F and G) Sequence motifs enriched in F) worst 10% and G) best 10% of 5′ UTRs by log2(RRS) from DART. H and I) Deletions of cap-proximal nucleotides significantly affect ribosome recruitment. 5′ UTRs were binned by the position of deleted nucleotides. Deletions of nucleotides 1 through 6 or 7 through 12 H) were more likely to cause a significant change (Benjamini-Hochberg adjusted p < 0.01) in ribosome recruitment and I) caused larger magnitudes of change in ribosome recruitment by DART. J) Gain of N1-methylpseudouridines and loss of guanosine nucleotides within the first 6 nucleotides increases ribosome recruitment in a dose-dependent manner. 5′ UTRs from the scanning deletion library were binned based on the change in the number of each nucleotide. Change in RRS relative to the respective parent sequences is plotted on the y-axis. K) N1-methylpseudouridines or guanosines are enriched in highly active or minimally active 5′ UTRs, respectively. UTRs were binned based on the top 10% (solid lines) or bottom 10% (dashed lines) by ribosome recruitment scores. Plot displays the percent of UTRs containing N1-methylpseudouridine (top) or guanosine (bottom) at each position.

## Human DART Protocol

### Buffers

- DNA Elution Buffer: 300 mM NaCl, 10 mM Tris, pH 8.0
- RNA Elution Buffer: 300 mM NaOAc pH 5.3, 1 mM EDTA pH 8.0
- 10x Translation Mix: 160 mM HEPES KOH pH 7.5, 1 mM spermidine, 8 mM ATP, 1 mM GTP, 400 mM potassium glutamate, 20 mM Mg glutamate, 200 mM creatine phosphate
- RNA Binding Buffer: 8M guanidine hydrochloride, 20mM EDTA, 20mM MES

### Pool amplification and purification

1. Prepare the pool PCR master mix (11 reactions per experiment):

2. Aliquot the master mix into PCR tubes, 50 µL per tube.
3. Run the PCR:

| Reagent | 1X | 11X |
| --- | --- | --- |
| 5X HF Buffer | 10 μL | 110 μL |
| oCL01 T7 Fwd primer (10uM) | 1 μL | 11 μL |
| oCL_Bar1 to 8 Barcoded Rev Primer (10uM) | 1 μL | 11 μL |
| dNTPs (10mM) | 1 μL | 11 μL |
| DMSO | 1.5 μL | 16.5 μL |
| 5' UTR pool (0.5ng/uL) | 1 μL | 11 μL |
| ddH2O | 34 μL | 374 μL |
| Phusion polymerase | 0.5 μL | 5.5 μL |
| **Total** | **50 μL** | **550 μL** |

4. Run each across 2 8% non-denaturing 1X TBE-polyacrylamide gel (run @ 200V for ∼45 minutes in 0.5X TBE).
5. Transfer the gel into a container to stain.
6. Add 20 mL of 0.5X TBE and 2 µL of SYBR gold per gel, stain at room temp on rocker for 5 minutes.
7. Visualize under blue light and excise amplified pool.
8. Transfer the gel slices to two 1.5 mL tubes containing 750 µL of DNA elution buffer.
9. Rock overnight at room temperature to elute DNA.
10. Transfer the eluted DNA solution to a new 1.5mL tube.
11. To precipitate, add 750 µL (1 volume) of isopropanol and 2 µL of Glycoblue. Incubate at - 20C for at least 30 minutes. (Potential pause point: DNA can be left in isopropanol until ready for next steps)
12. Spin at maximum speed in microcentrifuge for 30 minutes at 4C to pellet DNA.
13. Wash pellets with 750 µL of 70% ethanol.
14. Spin down at maximum speed for 10 minutes at 4C.
15. Resuspend the DNA pellets in a total of 8 µL of water.

2. Aliquot the master mix into PCR tubes, 50 μL per tube. 3. Run the PCR:
| Step | Temp | Time | Cycles |
| --- | --- | --- | --- |
| Initial denaturation | 98C | 30 sec | 1 |
| Denature | 98C | 10 sec | 18 |
| Anneal | 58C | 30 sec | 18 |
| Extend | 72C | 30 sec | 18 |
| Final extension | 72C | 5 min | 1 |
| Hold | 4C | Infinite | N/A |

### In vitro transcribe pool RNA

#### Uncapped RNA for enzymatic capping

1. Set up the in vitro transcription reactions:

| **Reagent** | **Amount** |
| --- | --- |
| 10X Reaction Buffer | 2 µL |
| ATP (75mM) | 2 µL |
| CTP (75mM) | 2 µL |
| GTP (75mM) | 2 µL |
| UTP (75mM) | 2 µL |
| Pool Template | 8 µL |
| T7 Enzyme Mix | 2 µL |
| **Total** | **20 µL** |

\*UTP was substituted with an equal concentration of N1-methylpsedouridine triphosphate for modified RNA DART analysis.

#### Co-transcriptionally Capped RNA

1. Set up the in vitro transcription reactions:

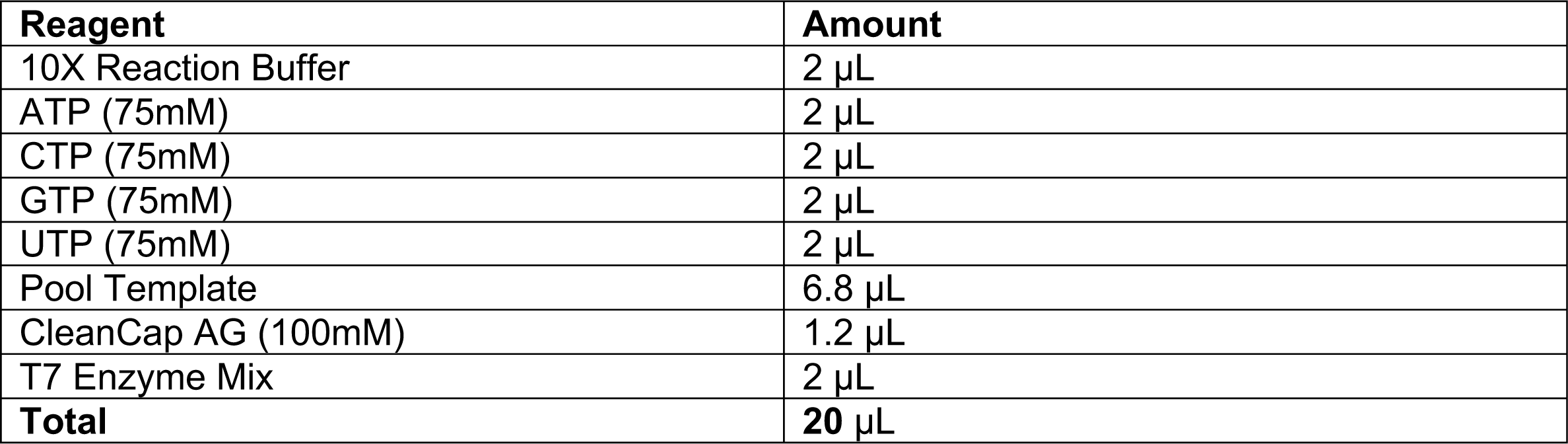
2. Incubate the transcription reaction at 37C for 2 hours
3. Add another 2 µL of T7 enzyme mix
4. Incubate the reaction at 37C overnight
5. Add 2 µL of TURBO DNase to the reaction to degrade template DNA. Incubate at 37C for 15 minutes.
6. Cast one 8% denaturing TBE-urea-polyacrylamide mini gel per reaction. Pre-run gels in 0.5X TBE for 20 minutes at 200V.
7. While gels are pre-running, add 24 µL of 2X formamide loading buffer to the RNA samples and denature RNA at 65C for 5 minutes.
8. Flush the wells of the gel with a syringe to remove urea from the wells.
9. Load 6 µL of the denatured RNA sample to each well of the gel. Run at 200V for ∼1 hour.
10. Transfer the gels to containers to stain.
11. Add 20 mL of 0.5X TBE and 2uL of SYBR gold, stain at room temp on rocker for 5 minutes.
12. Visualize under blue light and excise the RNA bands.
13. Transfer gel slices to tubes containing 750 µL of RNA elution buffer.
14. Elute overnight at 4C on rotator.
15. Transfer eluted RNA to a new 1.5mL tube.
16. To precipitate, add 750 µL of isopropanol (1 volume) and 2 µL of Glycoblue. Incubate at - 20C for at least 30 minutes. (Potential pause point, as before)
17. Spin at maximum speed for 30 minutes at 4C to pellet the RNA.
18. Wash pellets with 750 µL of 80% ethanol.
19. Spin down at maximum speed for 10 minutes at 4C.
20. Resuspend in 50uL of water for RNA going into capping, determine concentration and total yield with nanodrop. Store at −80 C.

**Co-transcriptionally Capped RNA** 1. Set up the in vitro transcription reactions:
| **Reagent** | **Amount** |
| --- | --- |
| 10X Reaction Buffer | 2 µL |
| ATP (75mM) | 2 µL |
| CTP (75mM) | 2 µL |
| GTP (75mM) | 2 µL |
| UTP (75mM) | 2 µL |
| Pool Template | 6.8 µL |
| CleanCap AG (100mM) | 1.2 µL |
| T7 Enzyme Mix | 2 µL |
| **Total** | **20 µL** |

#### Adding the 5’ m7G cap

1. Mix 50uL of gel-purified RNA with 2 µL of RNasin Plus. Incubate at 65C for 5 minutes to denature. Transfer to ice for 2 minutes.
2. Add the NEB Vaccinia capping system reagents and mix:

3. Incubate at 37C for 1 hour.

| Reagent | Amount |
| --- | --- |
| 10X Capping Buffer | 8 μL |
| GTP (10mM) | 12 μL |
| SAM (32mM) | 4 μL |
| Capping enzyme | 4 μL |
| **Total final volume** | **80 μL** |

#### Cleaning up the capping reaction (Zymo V column cleanup)

1. Add 560 µL of binding buffer to the capping reaction.
2. Add 640 µL of isopropanol and mix.
3. Place Zymo V columns in a collection tube. Apply 600 µL of RNA solution to the column.
4. Spin at maximum speed for 1 minute at 4C. Discard flow-through.
5. Wash column twice with 200 µL of 80% ethanol (same speed and time as above). Discard the flow-through.
6. Spin column once more to remove residual ethanol.
7. Transfer the column to a clean 1.5 mL tube. Add 200 µL of H2O to the resin.
8. Elute RNA by spinning at maximum speed for 2 minutes at 4C.
9. Add 22.2 µL of 3M sodium acetate, pH 5.3 and 1mL of 100% ethanol to precipitate RNA. Incubate at −20C for at least 30 minutes.
10. Spin at maximum speed at 4C for 30 minutes to pellet RNA.
11. Wash RNA with 80% ethanol, spin for another 10 minutes at max speed, 4C.
12. Resuspend RNA pellet in H2O, nanodrop to determine concentration. Record the yield.

#### Sucrose gradient preparation

-Before you begin, place the SW41 rotor and buckets in the cold room to cool.

1. Prepare a fresh stock of 10 mg/mL cycloheximide. Keep on ice.
2. Prepare the 10% and 50% sucrose solutions:

3. Mark the halfway point of 6 SW41 tubes with the gradient master tool (lower notch).
4. Add 10% sucrose to the SW41 tubes until ∼2-3mm above the halfway mark.
5. Draw 8 mL of 50% sucrose into a 10 mL syringe with a wide-gauge needle.
6. Gently depress plunger until sucrose solution is at the end of the needle. Keep tension on the plunger to prevent dripping.
7. Insert the needle into the SW41 tube containing the 10% sucrose to the bottom of the tube.
8. Gently depress the plunger to expel the 50% sucrose solution until the interface reaches the halfway mark on the tube.
9. Apply tension to the plunger to prevent dripping and slowly withdraw the syringe until the tip of the needle is slightly below the interface.
10. Quickly withdraw the needle through the 10% sucrose solution and out of the tube. Repeat with the remaining tubes.
11. Gently apply the SW41 long caps to the gradient tubes, allowing any air bubbles to vent through the hole in the cap.
12. Remove any extra liquid that may have accumulated on top of the cap.
13. Level the gradient station and insert tubes into the magnetic holder. Gently attach the holder to the gradient station.
14. Run the SW41_10-50%_Long program on the gradient master to establish the sucrose gradients.
15. Once finished, transfer the tubes into pre-chilled SW41 buckets. Place in cold room for at least 30 minutes to chill the gradients.

| Reagent | 10% amount | 50% amount |
| --- | --- | --- |
| HEPES KOH pH 7.4 (1M) | 1.2 mL | 1.2 mL |
| Magnesium glutamate (1M) | 120 μL | 120 μL |
| Potassium glutamate (1M) | 6 mL | 6 mL |
| Triton X-100 | 600 μL | 600 μL |
| DTT (1M) | 180 μL | 180 μL |
| Cycloheximide (10 mg/mL) | 600 μL | 600 μL |
| Sucrose (2M) | 8.76 mL | 43.8 mL |
| H<sub>2</sub>O | 42.54 mL | 7.5 mL |
| **Total** | **60 mL** | **60 mL** |

#### In vitro translation

1. Dilute RNA samples in water to 40 pmol for capped in 95 µL total volume in 1.5 mL tubes.
2. Thaw the HeLa lysate and clarify by centrifugation at 12,000 x g, for 1 minute at 4C.
3. Prepare the in vitro translation reaction master mix (7X).

4. Mix well and add 405 uL of the translation master mix to tubes containing 95 µL of pool RNA.
5. Incubate at 37C, 600 RPM on thermomixer for 30 minutes.
6. Reserve 50 µL of the translation reaction as input. Snap freeze and store at −80C.
7. Immediately proceed to fractionation.

| Reagent | 1X amount | 7X amount |
| --- | --- | --- |
| HeLa extract | 250 $\mu\text{L}$ | 1750 $\mu\text{L}$ |
| Cycloheximide (10 mg/mL) | 25 $\mu\text{L}$ | 175 $\mu\text{L}$ |
| RNasin Plus | 3.5 $\mu\text{L}$ | 24.5 $\mu\text{L}$ |
| cOmplete Mini EDTA-Free<br>Protease inhibitor (1 tablet in<br>2 mL H<sub>2</sub>O) | 17 $\mu\text{L}$ | 119 $\mu\text{L}$ |
| Translation Mix (10X) | 50 $\mu\text{L}$ | 350 $\mu\text{L}$ |
| H<sub>2</sub>O | 31.4 $\mu\text{L}$ | 219.8 $\mu\text{L}$ |
| CPK (10 mg/mL) | 16.5 $\mu\text{L}$ | 115.5 $\mu\text{L}$ |
| DTT (0.5M) | 1.6 $\mu\text{L}$ | 11.2 $\mu\text{L}$ |
| PMSF (0.1M) | 10 $\mu\text{L}$ | 70 $\mu\text{L}$ |
| **Total** | **405 $\mu\text{L}$** | **2835 $\mu\text{L}$** |

#### Sucrose gradient fractionation

1. Turn on the ultracentrifuge and allow to cool to 4C.
2. Load the remaining 450 µL of the translation reaction onto the top of the sucrose gradient by slowly dispensing against the side of the tube.
3. Balance the buckets + caps to within 0.1g by adding the 10% sucrose solution where necessary.
4. Screw on the caps to the SW41 buckets and attach to the rotor. Insert the rotor into the ultracentrifuge.
5. Centrifuge at 35,000 RPM for 3 hours at 4C
6. Fractionate the gradients on the gradient master. Pool the fractions corresponding to the 80S peak in a 15 mL conical tube.
7. Snap freeze samples and store in −80 C or proceed directly to hot phenol extraction

#### 80S hot phenol RNA extraction

1. Add 650 µL of acid phenol and 40 µL of 20% SDS per 600 uL of aqueous volume.
2. Prepare a 65C water bath by microwaving 1.5L of water for ∼5 minutes. Transfer to a 4L beaker. Start a second 1.5L in the microwave.
3. While the second bath is heating, place RNA-phenol tubes into the bath. Incubate for 5 minutes, vortexing every minute.
4. Transfer the samples to the now-heated second water bath. Repeat the incubation/vortexing for another 5 minutes.
5. Cool the samples for 5 minutes on ice.
6. Add a volume of chloroform equal to the acid phenol volume to pre-spun 15mL MaXtract tubes.
7. Transfer samples to tubes containing chloroform. Mix well by inversion.
8. Spin at 1500 x g for 5 minutes at 4C to separate the phases.
9. Transfer the aqueous fraction to new 15mL tubes.
10. Add 650 µL of acidic PCA per 600 µL of aqueous volume. Mix well by inversion.
11. Spin at maximum speed for 5 minutes at 4C to separate the phases.
12. Transfer the aqueous fraction to a new 15mL tube.
13. Add 650 µL of chloroform per 600 µL of aqueos volume. Mix well by inversion
14. Spin at maximum speed for 5 min at 4C to separate phases
15. Separate into 800 µL aliquots in 2 mL tubes. Add 89 µL of 3M NaOAc, 2 µL of glycoblue, and 1 mL of 100% isopropanol to each tube.
16. Incubate at −20C for at least 30 minutes.
17. Spin at maximum speed at 4C for 30 minutes to pellet RNA.
18. Wash RNA with 80% ethanol, spin for another 10 minutes at max speed, 4C. Repeat this for a second wash.
19. Store RNA in ethanol until ready for reverse transcription.

#### Reverse Transcription

1. Resuspend the RNA from one replica in 11 µL of water.
2. Add 1 µL of 10uM oCL04 RT primer.
3. Heat at 65C for 5 minutes, then step down by 5C every 2 minutes to 45C to anneal the primer (on thermocycler).
4. Add a master mix containing the following (per reaction):

4 µL 5X First-Strand Buffer
1 µL 0.1M DTT
1 µL 10mM dNTPs
1 µL RNasin Plus
1 µL Superscript III enzyme
5. Mix and incubate at 50C for 1 hour.
6. Add 2 µL of 1M NaOH, mix, incubate at 98C for 30 minutes to degrade RNA.
7. Neutralize with equivalent volume of 1M HCl.
8. Add 24 µL of 2X formamide loading buffer, denature at 80C for 2 minutes.
9. Run the samples on an 8% denaturing gel (each sample will be two lanes).
10. Cut bands, elute overnight, store in isopropanol or ethanol.

#### 5’ Adaptor Ligation

1. Resuspend cDNA in 5 µL of 10mM Tris-HCl pH 7.5.
2. Add 0.8 µL of 80uM oCL03 5’ Adaptor and 1 µL of 100% DMSO.
3. Heat at 75C for 2 minutes, place on ice.
4. Prepare the ligation master mix (per reaction):

- 2 µL of 10X RNA Ligase Buffer
- 0.2 µL of 100mM ATP
- 9 µL of 50% PEG 8000 (pre-warm this to 37C to make it easier to pipet)
- 1.1 µL of H20
- 1 µL of High Concentration T4 RNA Ligase
5. Add 13.3 µL of the master mix to cDNA-adapter solution
6. Mix, incubate at room temp overnight, 1000RPM on thermomixer

#### Adaptor Ligation Clean Up – Silane Beads

1. Vortex to resuspend the beads. Transfer 10 µL of bead slurry to a LoBind 1.5mL tube.
2. Magnetically separate, remove supernatant.
3. Wash beads once with 500 µL of RLT, separate, discard supernatant.
4. Resuspend beads in 60 µL of RLT and add to ligation reaction.
5. Add 60 µL of 100% EtOH
6. Mix by pipetting (leave tip in the tubes), incubate for 5 minutes at room temp mixing twice more.
7. Magnetically separate, remove supernatant, add 1mL of 75% EtOH.
8. Transfer sample resuspended in 75% EtOH to a new LoBind tube.
9. Incubate 30 seconds, separate, remove supernatant. Wash twice more with 75% EtOH.
10. Spin down briefly and remove residual EtOH with a 20 µL tip.
11. Air dry for 5 minutes
12. Resuspend in 27 µL of 10mM Tris-HCl pH 7.5 to elute.
13. Incubate for 5 minutes, separate, transfer eluted DNA to a new tube.

#### Diagnostic Library PCR Amplification

PCR amplify from adaptor-ligated cDNA at varying cycle numbers (6, 9, 12, 15, 18) in 16 µL reactions to determine optimal conditions for amplification:

| Reagent | 1X amount | 5x amount |
| --- | --- | --- |
| 5X HF Buffer | 3.2 μL | 16 μL |
| Fwd primer (i5XX) | 0.32 μL | 1.6 μL |
| Rev primer (i7XX) | 0.32 μL | 1.6 μL |
| dNTPs (10mM) | 0.32 μL | 1.6 μL |
| DMSO | 0.48 μL | 2.4 μL |
| DNA | 0.32 μL | 1.6 μL |
| ddH<sub>2</sub>O | 10.88 μL | 54.4 μL |
| Phusion polymerase | 0.16 μL | 0.8 μL |
| **Total** | **16 μL** | **80 μL** |

| Step | Temp | Time | Cycles |
| --- | --- | --- | --- |
| Initial denaturation | 98C | 30 sec | 1 |
| Denature | 98C | 10 sec | XX |
| Anneal | 72C | 15 sec | XX |
| Extend | 72C | 20 sec | XX |
| Final extension | 72C | 5 min | 1 |
| Hold | 4C | Infinite | N/A |

#### Final Library PCR Amplification

After determining optimal cycling conditions, scale up to 100 µL PCR reactions (in 50 µL aliquots):

Final Library PCR Amplification After determining optimal cycling conditions, scale up to 100 μL PCR reactions (in 50 μL aliquots):
| Reagent | 1x amount |
| --- | --- |
| 5X HF Buffer | 20 μL |
| i5 Fwd primer (10 μM) | 2 μL |
| i7 Rev primer (10 μM) | 2 μL |
| dNTPs (10mM) | 2 μL |
| DMSO | 3 μL |
| DNA | 2 μL |
| ddH<sub>2</sub>O | 68 μL |
| Phusion polymerase | 1 μL |
| **Total** | **100 μL** |

-Clean up and concentrate final libraries using SPRI bead clean up.

#### SPRI cleanup library

1. Resuspend settled AmpureXP beads well by vortexing.
2. Add 180 µL of bead suspension (do not separate) per 100 µL PCR reaction, pipette mix well, incubate at room temperature for 10 minutes. Pipette mix three times during the incubation.
3. Magnetically separate, wash the beads twice with 75% EtOH.
4. Air dry the beads for 5 minutes on magnet.
5. Remove from magnet, resuspend beads in 20 uL H2O, incubate for 5 minutes at room temperature.
6. Magnetically separate, transfer 18 µL of eluted library to new 1.5 mL tubes.
7. Add 6uL 6X Loading Dye to the samples.
8. Run libraries on 8% non-denaturing polyacrylamide gel.
9. Excise final library bands, elute in 750 µL DNA elution buffer overnight at room temperature.
10. Measure the library concentration using the Qubit High Sensitivity DNA assay.
11. Pool libraries to make a 1-10 nM final library.
12. Proceed to Illumina 2×150 sequencing.

